## Supplementary material for "Gene regulatory network reconstruction using single-cell RNA sequencing of barcoded genotypes in diverse environments": Jackson_2019_Supplemental_Data_1.tar.gz: data_imputation.R

```
MAGIC.FILE <- "MAGIC_DATA.tsv.gz"
VIPER.FILE <- "VIPER_DATA.tsv.gz"
SCIMPUTE.FILE <- "SCIMPUTE_DATA.tsv.gz"
SCTRANSFORM.FILE <- "SCTRANSFORM_DATA.tsv.gz"

if (!exists('count_data')) {source('figure_common_functions.R')}

require(Rmagic)
magic_raw <- dplyr::select(count_data, -META.DATA.COLUMNS) # Extract the UMI count data
magic_raw <- magic_raw[, base::colSums(magic_raw) > 0] # Select genes that are not all 0
magic_raw <- Rmagic::library.size.normalize() # Do whatever MAGIC wants for library size normalizing 
magic_raw <- sqrt(magic_raw) # Do square root instead of log2 x+1
magic_imputation <- Rmagic::magic(magic_raw, genes="all_genes", n.jobs = MP.CORES)
magic_imputation <- base::cbind(as.data.frame(magic_imputation$result), dplyr::select(count_data, META.DATA.COLUMNS))
file.connection <- gzfile(file.path(DATA.PATH, MAGIC.FILE), "w")
write.table(magic_imputation, file.connection, quote=FALSE, sep="\t")
close(file.connection)
magic_raw <- NULL
magic_imputation <- NULL

require(VIPER)
viper_raw <- dplyr::select(count_data, -META.DATA.COLUMNS) # Extract the UMI count data
viper_raw <- viper_raw[, base::colSums(viper_raw) > 0] # Select genes that are not all 0
viper_results <- VIPER::VIPER(viper_raw, num = 1000, percentage.cutoff = 0.5, minbool = FALSE, alpha = 0.5, report = FALSE, outdir = NULL, prefix = NULL)
viper_results <- base::cbind(as.data.frame(viper_results$imputed), dplyr::select(count_data, META.DATA.COLUMNS))
file.connection <- gzfile(file.path(DATA.PATH, VIPER.FILE), "w")
write.table(viper_results, file.connection, quote=FALSE, sep="\t")
close(file.connection)
viper_raw <- NULL
viper_results <- NULL

require(scImpute)
scimpute_raw <- dplyr::select(count_data, -META.DATA.COLUMNS)
scimpute_raw <- scimpute_raw[, base::colSums(scimpute_raw) > 0] # Select genes that are not all 0
condition.labels <- base::as.numeric(base::as.factor(count_data$Condition))
scimpute_raw <- t(scimpute_raw)

# This is from ScImpute's read_counts() function
scimpute_raw <- base::sweep(scimpute_raw, MARGIN = 2, 10^6/base::colSums(scimpute_raw), FUN = "*") 
scimpute_raw <- log10(scimpute_raw + 1.01)

# Call scImpute's handler directly instead of going through the wrapper that requires a file
# This still spits out RDS files though
# I cannot be bothered to fix that. Just delete them by hand.

scimpute_results <- scImpute:::imputation_wlabel_model8(count = scimpute_raw, labeled = TRUE, 
                                                        cell_labels = condition.labels, point = log10(1.01), 
                                                        drop_thre = 0.5, Kcluster = NULL, out_dir = DATA.PATH, 
                                                        ncores = 2)

# Do more stuff from ScImpute to match the output from the file wrapper
scimpute_data <- 10^(scimpute_results$count_imp - 1.01)
scimpute_data <- base::sweep(scimpute_data, MARGIN = 2, base::rowSums(dplyr::select(count_data, -META.DATA.COLUMNS)) / 10e6, FUN = "*")
scimpute_data <- base::cbind(as.data.frame(t(scimpute_data)), dplyr::select(count_data, META.DATA.COLUMNS))

file.connection <- gzfile(file.path(DATA.PATH, SCIMPUTE.FILE), "w")
write.table(scimpute_data, file.connection, quote=FALSE, sep="\t")
close(file.connection)
scimpute_raw <- NULL
scimpute_results <- NULL
scimpute_data <- NULL

require(sctransform)
vst_raw <- dplyr::select(count_data, -META.DATA.COLUMNS) # Extract the UMI count data
vst_model <- sctransform::vst(t(as.matrix(vst_raw)))
vst_denoised <- sctransform::denoise(vst_model)
```
