## Supplementary material for "Gene regulatory network reconstruction using single-cell RNA sequencing of barcoded genotypes in diverse environments": Jackson_2019_Supplemental_Data_1.tar.gz: figure_2.R

```
# Output paths
ENO2.RIDGELINE <- "figure_2a_eno2.png"
ADH2.RIDGELINE <- "figure_2a_adh2.png"
PDC1.RIDGELINE <- "figure_2a_pdc1.png"
FIGURE2A <- "figure_2a.png"
FIGURE2B <- "figure_2b.png"
FIGURE2 <- "figure_2.png"

if (!exists('count_data')) {source('figure_common_functions.R')}
if (!exists('sce.data')) {sce.data <<- dimred_for_umap(load_data_into_sce(count.data = dplyr::select(count_data, -META.DATA.COLUMNS), 
                                                                          meta.data = meta_data), 
                                                       random_state=RANDOM.SEED)}

# Select data for genes to include in figure 2a and melt it
ridgeline.genes <- c('YHR174W', 'YLR044C', 'YMR303C')
ridgeline.data <- count_data[, c(ridgeline.genes, 'Condition', 'Genotype_Group')]
ridgeline.data <- reshape2::melt(ridgeline.data, id.vars = c('Condition', 'Genotype_Group'), variable.name = "Gene", value.name="Count")

# Calculate the median expression values for each strain (TF deletion)
ridgeline.medians <- dplyr::group_by(ridgeline.data, Gene, Condition, Genotype_Group)
ridgeline.medians <- dplyr::summarize(ridgeline.medians, gene_median=median(Count))

# Plot function with the common options for each plot
make.ridgeline.plt <- function(ridge.data, median.data, gene.name) {
  ggplot(ridge.data[ridge.data$Gene == gene.name, ], aes(x=Count, y=Condition)) +
    stat_binline(aes(height=..ndensity.., fill=factor(Condition)), binwidth=1, center=0, draw_baseline = FALSE, scale = 0.9) + 
    geom_point(data=median.data[median.data$Gene == gene.name, ], aes(x=gene_median, y=Condition)) +
    theme_ridges(grid = FALSE) +
    theme(axis.text.x = element_text(size = 10, angle=90, vjust = 0.5, hjust = 0.5), axis.title.x = element_text(size = 12, face="bold", hjust=0.5),
          axis.text.y = element_text(size = 10, vjust=0.5, hjust = 1), axis.title.y = element_text(size = 12, face="bold", hjust=0.5),
          plot.title = element_text(size = 12, face = "bold.italic", hjust=0.5)) +
    scale_y_discrete(limits=CONDITION.LEVELS, labels=CONDITION.LABELS) +
    scale_fill_manual(values=CONDITION.COLORS) +
    theme(axis.text.y = element_blank(), axis.title.y = element_blank(), 
          axis.title.x = element_blank(),
          legend.position = 'none',
          plot.margin = unit(c(0, 0, 0, 0), "cm"))
}

eno2.plt <- make.ridgeline.plt(ridgeline.data, ridgeline.medians, 'YHR174W') +
  labs(title="ENO2") +
  scale_x_continuous(limits=c(-0.5,60), breaks=c(0, 20, 40))

pdc1.plt <- make.ridgeline.plt(ridgeline.data, ridgeline.medians, 'YLR044C') +
  labs(title="PDC1") +
  scale_x_continuous(limits=c(-0.5,60), breaks=c(0, 20, 40))

adh1.plt <- make.ridgeline.plt(ridgeline.data, ridgeline.medians, 'YMR303C') +
  labs(title="ADH2") +
  scale_x_continuous(limits=c(-0.5,21), breaks=c(0, 20))

plfig2a <- plot_grid(eno2.plt, pdc1.plt, adh1.plt, ncol = 3, align = 'v', labels = c("A)", "",""))#ggsave(FIGURE2A, plot=plfig2a, path=OUT.PATH, device="png", width=2, height=5.5, units="in", limitsize = FALSE)

# Figure 2B
print("Constructing Figure 2B")

p1umap.cond <- umap.plot(sce.data, color.by='Condition', alpha=0.5, size=0.05, color.title='Condition') + 
  scale_color_manual(labels = CONDITION.LABELS, breaks = CONDITION.LEVELS, values = CONDITION.COLORS) + 
  guides(colour = guide_legend(override.aes = list(size=4, alpha=1), reverse = TRUE)) +
  labs(title = "Growth Condition") +
  theme(legend.position = "none", axis.title = element_blank(), axis.text = element_blank(),
        plot.title = element_text(size=12, face = "bold", hjust = 0.5))

plumap.geno <- umap.plot(sce.data, color.by='Genotype_Group', alpha=0.5, size=0.05, color.title='Condition') + 
  scale_color_manual(labels = GENOTYPE.LABELS, breaks = GENOTYPE.LEVELS, values = GENOTYPE.COLORS) + 
  guides(colour = guide_legend(override.aes = list(size=4, alpha=1), reverse = TRUE)) +
  labs(title = "Strain Genotype") +
  theme(legend.position = "none", axis.title = element_blank(), axis.text = element_blank(),
        plot.title = element_text(size=12, face = "bold", hjust = 0.5))

figure.2b.plt <- plot_grid(p1umap.cond, plumap.geno, ncol = 1, align = 'v', labels = c("B)", "C)"), axis="lr")
#ggsave(plot=plfig2b, filename=FIGURE2B, path=OUT.PATH, device="png", width=21, height=8, units="in", limitsize = FALSE)

figure.2.plt <- plot_grid(plfig2a, figure.2b.plt, ncol = 2, align = "h", rel_widths = c(0.8, 1))
ggsave(plot=figure.2.plt, filename=FIGURE2, path=OUT.PATH, device="png", width=5, height=5, units="in", dpi = FIGURE.DPI)
```
