## Supplementary material for "Gene regulatory network reconstruction using single-cell RNA sequencing of barcoded genotypes in diverse environments": Jackson_2019_Supplemental_Data_1.tar.gz: figure_3.R

```
# Output paths
FIGURE3A.ORIENTED.COLUMNS <- FALSE
FIGURE3A <- 'figure_3a.png'
FIGURE3B <- 'figure_3b.pdf'
FIGURE3B.GENO <- 'figure_3b_genotype.pdf'
FIGURE3B.EXPR <- 'figure_3b_expression.pdf'
SUPPLEMENTAL.FIGURE.3A <- 'supplemental_figure_3A.png'
SUPPLEMENTAL.FIGURE.3B <- 'supplemental_figure_3B.pdf'

if (!exists('count_data')) {source('figure_common_functions.R')}

print("Loading Figure 3 Data")
sce.obj.fig3 <- load_data_into_sce(count.data = dplyr::select(count_data, -META.DATA.COLUMNS), meta.data = meta_data)
sce.conditions <- list()
for (condition in rev(CONDITION.LEVELS)) {sce.conditions[[condition]] <- sce.obj.fig3[, sce.obj.fig3$Condition == condition]}
rm(sce.obj.fig3)

print("Normalizing Figure 3 Data")
sce.conditions <- lapply(sce.conditions, normalizeSCE)
print("PCA for Figure 3 Data")
sce.conditions <- lapply(sce.conditions, function(x) {runPCA(x, ncomponents=50, ntop=Inf, method='irlba', exprs_values='logcounts')})
print("UMAP for Figure 3 Data")
sce.conditions <- lapply(sce.conditions, function(x) {runUMAP(x, random_state=RANDOM.SEED, min_dist=0.1, use_dimred='PCA')})
print("SNN for Figure 3 Data")
sce.conditions <- lapply(sce.conditions, apply.snn.cluster)

print("Generating Figure 3A")
genotype.plts <- lapply(names(sce.conditions), 
                        function(x) {umap.plot(sce.conditions[[x]], color.by='Genotype_Group', alpha=0.8, size=0.25, color.title='Genotype') + 
                            scale_color_manual(labels = GENOTYPE.LABELS, breaks = GENOTYPE.LEVELS, values = GENOTYPE.COLORS) +
                            theme(legend.position='None')})

colors.clust <- brewer.pal(9, 'Set1')
cluster.plts <- lapply(names(sce.conditions), 
                       function(x) {umap.plot(sce.conditions[[x]], color.by='cluster', alpha=0.8, size=0.25, color.title='cluster') + 
                           scale_color_manual(values=colors.clust)})

pir1.plts <- lapply(names(sce.conditions), 
                       function(x) {umap.plot(sce.conditions[[x]], gene = "YKL164C", alpha=0.8, size=0.25, color.title='cluster') + 
                           scale_colour_viridis(option='B', limits = c(0,4), oob = squish)})

dse2.plts <- lapply(names(sce.conditions), 
                    function(x) {umap.plot(sce.conditions[[x]], gene = "YHR143W", alpha=0.8, size=0.25, color.title='cluster') + 
                        scale_colour_viridis(option='B', limits = c(0,4), oob = squish)})

htb.plts <- lapply(names(sce.conditions), 
                   function(x) {umap.plot(sce.conditions[[x]], gene = c("YDR224C", "YBL002W"), alpha=0.8, size=0.25, color.title='cluster') + 
                       scale_colour_viridis(option='B', limits = c(0,4), oob = squish)})

if (FIGURE3A.ORIENTED.COLUMNS) {
  # Rerack the lists so that they're printed column-wise (does plot_grid have an option for this I can't find?)
  all.plts <- c()
  for (i in 1:11) {all.plts <- c(all.plts, cluster.plts[i], genotype.plts[i], pir1.plts[i], dse2.plts[i], htb.plts[i])}
  plot.ncol <- 5
  plot.width <- 5
  plot.height <- 11
} else {
  all.plts <- c(pir1.plts, dse2.plts, htb.plts, genotype.plts, cluster.plts)
  plot.ncol <- 11
  plot.height <- 5
  plot.width <- 11
}

all.plts <- lapply(all.plts, function(x) {x + theme(axis.text = element_blank(), axis.title = element_blank(), plot.title = element_blank(), legend.position = 'none')})
figure3a.plts <- plot_grid(plotlist = all.plts, ncol = plot.ncol, align = 'hv')
ggsave(plot=figure3a.plts, filename=FIGURE3A, path=OUT.PATH, device="png", width=plot.width, height=plot.height, units="in", dpi = FIGURE.DPI)

print("Generating Figure 3B / Supplemental Figure 3B")

sum_log2 <- function(df) {log2(rowSums(2 ^ df))}

plt.fig.3b <- function(sce.conditions, include.conditions = NULL) {
  #Gene systemic names
  gene.systemics <- c("HTB", "YKL164C", "YHR143W", "YGL253W")
  # Translate to common names
  gene.names <-yeast.systemic.to.common(gene.systemics)
  if (is.null(include.conditions)) {include.conditions <- CONDITION.LEVELS}
  condition.plt.labels <- translate.condition.name(include.conditions)
  
  # Get clusters and expression data
  extract.expression.clusters.from.list(sce.conditions, c("YDR224C", "YKL164C", "YBL002W", "YHR143W", "YGL253W"), include.names = include.conditions) %>%
    dplyr::mutate(HTB=sum_log2(.[c("YDR224C", "YBL002W")])) %>%
    dplyr::select(-c("YDR224C", "YBL002W")) %>%
    dplyr::group_by(clusters, condition) %>%
    dplyr::summarize_all(mean) %>% 
    reshape2::melt(id.vars = c("clusters", "condition"), value.name = "logcount", variable.name = "Gene") -> expr.data
  
  # Plot expression data
  expr.plt <- ggplot(expr.data, aes(x=clusters, y=Gene, fill=logcount)) +
    geom_tile(colour="white",size=0.25) +
    scale_y_discrete(limits = rev(gene.systemics), labels = rev(gene.names)) +
    scale_fill_viridis(option='B', limits = c(0,4), oob = squish) + 
    facet_wrap(~condition, ncol=11, labeller = as_labeller(structure(condition.plt.labels, names=include.conditions))) +
    theme(strip.background = element_blank(), strip.text = element_text(size=10, face = "bold"))
  
  #ggsave(plot=expr.plt, filename=FIGURE3B.EXPR, path=OUT.PATH, device=cairo_pdf, width=8.5, height=5, units="in", limitsize = FALSE)
  
  # Calculate the proportion of genotypes per [cluster + condition]
  extract.genotype.count(sce.conditions, include.names = include.conditions) %>%
    dplyr::group_by(clusters, condition, genotype) %>%
    dplyr::summarize(count = n()) -> geno.count.data
  geno.count.data %>%
    dplyr::group_by(condition, clusters) %>%
    dplyr::summarize(total_count = sum(count)) %>%
    dplyr::right_join(geno.count.data, by=c("condition", "clusters")) -> geno.count.data
  geno.count.data$proportion <- geno.count.data$count / geno.count.data$total_count
  
  # Plot genotypes per cluster (faceted on condition)
  geno.plt <- ggplot(geno.count.data, aes(x=clusters, y=proportion, fill = genotype)) +
    geom_bar(position='stack', stat='identity') + 
    scale_fill_manual(breaks = GENOTYPE.LEVELS, values = GENOTYPE.COLORS) + 
    ylab('proportion') + xlab('') +
    theme(axis.text.x = element_text(angle = 90, hjust = 1, vjust=0.5)) +
    facet_wrap(~condition, ncol=11, labeller = as_labeller(structure(condition.plt.labels, names=include.conditions))) +
    theme(strip.background = element_blank(), strip.text = element_text(size=10, face = "bold"))
  
  #ggsave(plot=geno.plt, filename=FIGURE3B.GENO, path=OUT.PATH, device=cairo_pdf, width=8.5, height=5, units="in", limitsize = FALSE)
  
  plot.stack <- plot_grid(geno.plt, expr.plt, align = "v", rel_heights=c(1, 0.75), ncol=1)
  return(plot.stack)
}

fig.3b.conditions <- c("YPD", "YPDRapa")
supp.fig.3b.conditions <- CONDITION.LEVELS[!(CONDITION.LEVELS %in% fig.3b.conditions)]
ggsave(FIGURE3B, plot=plt.fig.3b(sce.conditions, fig.3b.conditions), path=OUT.PATH, device=cairo_pdf, width=6, height=4, units="in", dpi = FIGURE.DPI)
ggsave(SUPPLEMENTAL.FIGURE.3B, plot=plt.fig.3b(sce.conditions, supp.fig.3b.conditions), path=OUT.PATH, device=cairo_pdf, width=20, height=4, units="in", dpi = FIGURE.DPI)

print("Generating Supplemental Figure 3A")
cond.cluster.markers <- extract.markers(sce.conditions, marker.cutoff.num=10)

plt.list <- list()
rel.height <- c()
for (cond.name in names(sce.conditions)) {
  # Get the marker gene names for this condition
  cond.markers <- cond.cluster.markers[cond.cluster.markers$condition == cond.name, ]
  
  # Extract the marker gene expression data from this condition SCE
  cond.expr <- extract.expression.clusters(sce.conditions[[cond.name]], unique(cond.markers$Gene))
  cond.clusters <- as.numeric(cond.expr$clusters)
  cond.expr <- dplyr::select(cond.expr, -c("condition", "genotype", "clusters"))
  
  # Hclust within each cluster of cells to determine x-axis ordering
  cond.x.order <- lapply(1:max(as.numeric(cond.clusters)), function(x) {hclust(dist(cond.expr[cond.clusters == x,]), 'average')})
  cond.x.order <- lapply(cond.x.order, function(x) {as.character(x$labels[x$order])})
  cond.x.order <- unlist(cond.x.order)
  
  # Z-Score for plotting
  cond.expr <- as.data.frame(scale(cond.expr))
  
  # Melt for heatmap
  cond.expr$cluster <- cond.clusters
  cond.expr$cell <- rownames(cond.expr)
  cond.expr <- reshape2::melt(cond.expr, id.vars = c("cell", "cluster"), value.name = "expression", variable.name = "gene")
  cond.expr$cell <- factor(cond.expr$cell, levels=cond.x.order)
  
  # Find the breaks between the cluster markers
  breaks <- cumsum(table(as.numeric(cond.clusters))) + 0.5
  
  # Enforce cluster ordering of y-axis
  cond.systemic.names <- unique(cond.markers[order(cond.markers$cluster), ]$Gene)
  cond.common.names <- yeast.systemic.to.common(cond.systemic.names)
  
  # Plot the heatmap
  cond.heatmap <- ggplot(cond.expr, aes(x=cell, y=gene, fill=expression)) +
    geom_tile() + 
    scale_fill_gradient2(low='slateblue2', mid='black', high='yellow', limits=c(-3, 3), oob=squish) + 
    theme(axis.text.y = element_text(size=4), 
          axis.title.x = element_blank(), 
          axis.text.x = element_blank(), 
          axis.title.y = element_blank(), 
          axis.ticks.x = element_blank(),
          plot.title = element_text(size=10, hjust=0.5, face="bold"),
          legend.position = "none") + 
    scale_y_discrete(limit=rev(cond.systemic.names), labels=rev(cond.common.names)) +
    geom_vline(xintercept=breaks, color='white', size=0.5) +
    labs(title = translate.condition.name(cond.name))

  rel.height <- c(rel.height, length(cond.systemic.names))
  plt.list[[cond.name]] <- cond.heatmap
  cond.heatmap <- cond.heatmap + theme(axis.text.y = element_text(size=6))
  #ggsave(paste0(substr(SUPPLEMENTAL.FIGURE.3A, 1, 22), "_", cond.name, ".png"), plot = cond.heatmap, path=OUT.PATH, width = 8.5, height = 2 + length(cond.systemic.names) * 0.1, units = "in", device = "png")
}

rel.height <- rel.height / max(rel.height)
plt.grid <- plot_grid(plotlist = plt.list, align = "v", ncol=1, rel_heights = rel.height)
ggsave(SUPPLEMENTAL.FIGURE.3A, plot = plt.grid, path=OUT.PATH, width = 8.5, height = 48, units = "in", device = "png")
```
