## Supplementary material for "Gene regulatory network reconstruction using single-cell RNA sequencing of barcoded genotypes in diverse environments": Jackson_2019_Supplemental_Data_1.tar.gz: figure_4.R

```
require(DESeq2)

# Output paths
FIGURE.4.FILE <- 'figure_4.pdf'
FIGURE.4A.FILE <- 'figure_4a.png'
FIGURE.4B.FILE <- 'figure_4b.png'
SUPPLEMENTAL.FIGURE.4C.FILE <- 'supplemental_figure_4c.pdf'
FIGURE.4D.NETWORK.FILE <- 'figure_4d_network_file.tsv'

if (!exists('count_data')) {source('figure_common_functions.R')}
if (!exists('sce.data')) {sce.data <<- dimred_for_umap(load_data_into_sce(count.data = dplyr::select(count_data, -META.DATA.COLUMNS), 
                                                                          meta.data = meta_data), 
                                                       random_state=RANDOM.SEED)}

print("Creating Figure 4A / Supplemental Figure 4A")

fig.4a.conditions <- c("YPD", "AmmoniumSulfate", "YPDRapa", "Urea")
figure.4a.plt <- plt.gene.violins(sce.data, "YKR039W", conditions = fig.4a.conditions) + 
  facet_wrap(~condition, labeller = as_labeller(structure(CONDITION.LABELS, names=CONDITION.LEVELS)), ncol = 2) +
  labs(title="GAP1 Expression", x="Genotype", y="log2(Expression)")

#ggsave(FIGURE.4A.FILE, plot=figure.4a.plt, path=OUT.PATH, device="png", width=4.25, height=5, units="in")

print("Creating Figure 4B")
make.pseudobulk <- function(count.df) {
  count.df %>%
    dplyr::select(Genotype_Group, Genotype) %>%
    dplyr::group_by(Genotype) %>%
    dplyr::summarise_all(data.table::first) -> genotype.lookup
  
  count.df %>%
    dplyr::select(-c(Genotype_Group, tenXBarcode, Replicate)) %>%
    dplyr::group_by(Genotype, Condition) %>%
    dplyr::summarise_all(sum) %>% 
    dplyr::left_join(genotype.lookup, by="Genotype") %>%
    as.data.frame() -> bulked
  
  bulked$UMI <- base::rowSums(dplyr::select(bulked, -c(Genotype, Condition, Genotype_Group)))
  return(bulked)
}

print("Bulking data")
pseudo.bulk.counts <- make.pseudobulk(count_data)

# This is just to shut DESeq up about the perens
levels(pseudo.bulk.counts$Genotype_Group)[levels(pseudo.bulk.counts$Genotype_Group) == "WT(ho)"] <- "WT"

non.wt.genotypes <- GENOTYPE.LEVELS[GENOTYPE.LEVELS != "WT(ho)"]
non.wt.genotype.labels <- GENOTYPE.LABELS[GENOTYPE.LABELS != "WT"]

print("Analyzing data for differential gene expression [~Genotype]")
DESeq.results <- NULL
for (condition in CONDITION.LEVELS) {
  # Extract the data corresponding to a specific growth condition and pack it into a DESeq DataSet
  pb.condition <- dplyr::filter(pseudo.bulk.counts, Condition == condition)
  pb.condition.meta <- dplyr::select(pb.condition, c(Genotype_Group, Genotype, UMI))
  pb.condition <- dplyr::select(pb.condition, -c(Genotype_Group, Genotype, Condition, UMI))
  pbc.deseq <- DESeq2::DESeqDataSetFromMatrix(countData = t(pb.condition), 
                                              colData = pb.condition.meta, 
                                              design = ~Genotype_Group)
  pbc.deseq$Genotype_Group <- relevel(pbc.deseq$Genotype_Group, "WT")
  
  # Run DESeq
  pbc.deseq <- DESeq2::DESeq(pbc.deseq,
                             fitType = "local", 
                             quiet = TRUE, 
                             minReplicatesForReplace = Inf,
                             parallel = TRUE,
                             BPPARAM = BiocParallel::MulticoreParam(MP.CORES))
  
  # Pull out all of the WT vs TF deletion results and pack them into a data.frame
  for (genotype.name in non.wt.genotypes) {
    pbc.results <- DESeq2::results(pbc.deseq, 
                                   contrast = c("Genotype_Group", genotype.name, "WT"), 
                                   lfcThreshold = FIG4.LFC.THRESHOLD)
    if (base::nrow(pbc.results) == 0) {next}
    pbc.results$genotype <- genotype.name
    pbc.results$condition <- condition
    pbc.results$gene <- rownames(pbc.results)
    rownames(pbc.results) <- NULL
    pbc.results <- as.data.frame(pbc.results)
    DESeq.results <- base::rbind(DESeq.results, pbc.results)
  }
}

print("Generating Figure 4B")
DESeq.results.significant <- dplyr::filter(DESeq.results, padj < FIG4.P.ADJ.THRESHOLD)

# Summarize the differential gene expression results for plotting
DESeq.results.significant %>%
  dplyr::group_by(condition, genotype) %>%
  dplyr::summarise(DiffGenes = n()) %>%
  dplyr::right_join(base::expand.grid(condition = CONDITION.LEVELS, 
                                      genotype = non.wt.genotypes, 
                                      stringsAsFactors = FALSE), 
                    by=c("condition", "genotype")) %>%
  as.data.frame() -> figure.4b.data

figure.4b.plt <- ggplot(figure.4b.data, aes(x=condition, y=genotype, fill=DiffGenes)) +
  theme_classic() +
  geom_tile() +
  geom_text(aes(label=DiffGenes)) +
  scale_x_discrete(limits=rev(CONDITION.LEVELS), labels=rev(CONDITION.LABELS)) +
  scale_y_discrete(limits=non.wt.genotypes, labels=non.wt.genotype.labels) +
  scale_fill_gradient(low="white", high=muted("blue"), na.value = "white") +
  theme(axis.text.x = element_text(size = 10, angle = 90, vjust=0.5, hjust=1), axis.title.x = element_text(size=12),
        axis.text.y = element_text(size = 10), axis.title.y = element_text(size=12),
        plot.title = element_text(size = 12, face = "bold", hjust = 0.5),
        legend.position = "none") +
  labs(x = "Growth Condition", y = "Genotype", title = "Differential Expression vs WT [# Genes]")

#ggsave(FIGURE.4B.FILE, plot=figure.4b.plt, path=OUT.PATH, device="png", width=4.25, height=5, units="in")

print("Generating Figure 4C")
# Extract the gene log fold changes
supp.figure.4c.plts <- list()
for (cond in CONDITION.LEVELS) {
  # Get the significant results for this condition
  cond.data <- DESeq.results.significant[DESeq.results.significant$condition == cond, ]
  cond.data <- cond.data[base::toupper(cond.data$genotype) != yeast.systemic.to.common(cond.data$gene), ]
  if (base::nrow(cond.data) == 0) {next}
  
  # Extract all of the DESeq results for each significant gene (including the comparisons that aren't significant)
  DESeq.results %>%
    dplyr::filter(condition == cond) %>%
    dplyr::filter(gene %in% unique(cond.data$gene)) -> cond.results
  
  # Replace P.ADJ with an asterisk if it's significant and an NA if it isnt
  cond.results$padj[cond.results$padj >= FIG4.P.ADJ.THRESHOLD] <- NA
  cond.results$padj[cond.results$padj < FIG4.P.ADJ.THRESHOLD] <- "*"
  
  # Cluster the genes based on log2 fold-change and save the gene order as a vector
  pivoted.results <- reshape2::dcast(cond.results[, c("genotype", "gene", "log2FoldChange")], gene ~ genotype, value.var = "log2FoldChange")
  rownames(pivoted.results) <- pivoted.results$gene
  pivoted.results$gene <- NULL
  cond.x.order <- hclust(dist(pivoted.results), 'average')
  cond.x.order <- as.character(cond.x.order$labels[cond.x.order$order])
  
  # Draw the plot for this condition
  cond.plt <- ggplot(cond.results, aes(x=gene, y=genotype, fill=log2FoldChange)) +
    theme_classic() +
    geom_tile() +
    geom_text(aes(label=padj)) +
    scale_x_discrete(limits=cond.x.order, labels=yeast.systemic.to.common(cond.x.order)) +
    scale_y_discrete(limits=non.wt.genotypes, labels=non.wt.genotype.labels) +
    scale_fill_gradient2(high='brown1', mid='white', low='dodgerblue', limit=c(-2,2), oob=squish)  +
    theme(axis.text.x = element_text(size = 10, angle = 90, vjust=0.5, hjust=1), axis.title.x = element_text(size=12),
          axis.text.y = element_text(size = 10), axis.title.y = element_text(size=12),
          plot.title = element_text(size = 12, face = "bold", hjust = 0.5),
          legend.position = "none") +
    labs(x = "Gene", y = "Genotype", title = paste0("Fold Change (vs WT) [", translate.condition.name(cond), "]"))
  
  # If it's YPD, use this plot for Figure 4. Otherwise make it Supplemental Figure 4
  if (cond != "YPD") {
    supp.figure.4c.plts[[cond]] <- cond.plt + theme(axis.text.x = element_text(size = 4, angle = 90, vjust=0.5, hjust=1))
  } else {
    figure.4c.plt <- cond.plt + theme(axis.text.x = element_text(size = 6, angle = 90, vjust=0.5, hjust=1))
  }
  #ggsave(cond.plt + theme(legend.position = "right"), filename=cond.file, path=OUT.PATH, device = cairo_pdf, height=4.8, width=2.64+(0.18*length(cond.x.order)), units="in")
}

# Lay out the entire figure 4 grid
figure.4.grid <- plot_grid(plot_grid(figure.4a.plt, figure.4b.plt, align="h", labels=c("A)", "B)")), figure.4c.plt, ncol=1, labels=c("", "C)", rel_heights = c(1, 0.75)))
ggsave(FIGURE.4.FILE, plot=figure.4.grid, path=OUT.PATH, device=cairo_pdf, width=8.5, height=8, units="in", dpi = FIGURE.DPI)

# Lay out Supplemental Figure 4C
supp.figure.4c.plts <- plot_grid(plotlist = rev(supp.figure.4c.plts), ncol=1, align = "v")
ggsave(SUPPLEMENTAL.FIGURE.4C.FILE, plot=supp.figure.4c.plts, path=OUT.PATH, device=cairo_pdf, width=8.5, height=30, units="in", dpi = FIGURE.DPI)
```
