## Supplementary material for "Gene regulatory network reconstruction using single-cell RNA sequencing of barcoded genotypes in diverse environments": Jackson_2019_Supplemental_Data_1.tar.gz: figure_5.R

```
require(ggplot2)
require(dplyr)
require(reshape2)
require(cowplot)
require(stringr)

FIGURE.5.FILE <- 'figure_5.pdf'
FIGURE.5A <- 'figure_5a.pdf'
FIGURE.5B <- 'figure_5b.pdf'
FIGURE.5C <- 'figure_5c.pdf'
SUPPLEMENTAL.FIGURE.5A <- 'supplemental_figure_5a.pdf'

# File paths (set if they are not already in the global space)
if (!exists('count_data')) {source('figure_common_functions.R')}
if (!exists('STABLE5.FILE')) {STABLE5.FILE <- "STable5.tsv"}


setwd(WORKING.PATH)

STABLE5.DATA <- read.table(file.path(WORKING.PATH, DATA.PATH, STABLE5.FILE), sep = "\t", header=TRUE, stringsAsFactors = FALSE)

make_dotplot <- function(plot.data, x.var, y.var, x.lab=NULL, y.lab=NULL, plot.title=NULL, x.lim = c(NA, NA), y.lim = c(NA, NA), x.axis.log=TRUE) {
  pl <- ggplot(plot.data, aes_string(x=x.var, y=y.var, group=x.var)) +
    geom_jitter(color="gray", width = 0.1, alpha = 0.5) +
    coord_cartesian(ylim = y.lim) +
    labs(x = x.lab, y = y.lab, title = plot.title) +
    add_plot_theme()
  if (x.axis.log) {pl <- pl + scale_x_log10(breaks = c(1, 10, 100, 1000, 10000, 40000), limits = x.lim)}
  return(pl)
}

add_plot_theme <- function(pl) {
  theme_classic() +
    theme(panel.grid.major = element_blank(), panel.grid.minor = element_blank(),
          axis.text.x = element_text(size = 12, angle=90, vjust=0.5), axis.title.x = element_text(size = 12, face="bold", hjust=0.5),
          axis.text.y = element_text(size = 12), axis.title.y = element_text(size = 12, face="bold", hjust=0.5),
          plot.title = element_text(size = 12, face = "bold", hjust=0.5),
          strip.text = element_text(size = 12, face = "bold"),
          legend.position = "none")
}

# Figure 5A
print("Constructing Figure 5A")
fig.5a.data <- dplyr::filter(STABLE5.DATA, Figure == "Figure_5a")
fig.5a.levels <- c("Shuffled", "Neg Data", "No Impute", "MAGIC", "ScImpute", "VIPER")
fig.5a.labels <- c("Neg.(Shuffled)", "Neg. (Data)", "No Imputation", "MAGIC", "scImpute", "VIPER")

plimpute <- make_dotplot(fig.5a.data, 'Label', 'AUPR', NULL, 'AUPR', y.lim=c(0,max(fig.5a.data$AUPR)), x.axis.log = FALSE) +
  scale_x_discrete(limits=fig.5a.levels, labels = fig.5a.labels) +
  stat_summary(fun.data = mean_sdl, fun.args = list(mult = 1), color = "black",  position = position_nudge(x = 0.25, y = 0)) +
  theme(axis.text.x = element_blank())

plimpute2 <- make_dotplot(fig.5a.data, 'Label', 'N_PREC', NULL, '# Edges\n(Prec. > 0.5)', y.lim=c(0,max(fig.5a.data$N_PREC)), x.axis.log = FALSE) +
  scale_x_discrete(limits=fig.5a.levels, labels = fig.5a.labels) +
  stat_summary(fun.data = mean_sdl, fun.args = list(mult = 1), color = "black",  position = position_nudge(x = 0.25, y = 0)) +
  theme(axis.text.x = element_blank())

plimpute3 <- make_dotplot(fig.5a.data, 'Label', 'N_CONF', NULL, '# Edges\n(Conf. > 0.95)', y.lim=c(0,max(fig.5a.data$N_CONF)), x.axis.log = FALSE) +
  scale_x_discrete(limits=fig.5a.levels, labels = fig.5a.labels) +
  stat_summary(fun.data = mean_sdl, fun.args = list(mult = 1), color = "black",  position = position_nudge(x = 0.25, y = 0)) +
  theme(axis.text.x = element_blank())

plimpute4 <- ggplot(fig.5a.data, aes_string(x='Label', y='N_CONF', group='Label')) +
  geom_blank() +
  scale_x_discrete(limits = fig.5a.levels, labels = fig.5a.labels) +
  scale_y_continuous(limits = c(0,0)) +
  labs(x = "Imputation") +
  add_plot_theme() +
  theme(axis.line = element_blank(), axis.title.y = element_blank(), axis.text.y = element_blank(), 
        axis.ticks = element_blank(), axis.text.x = element_text(hjust=1))

impute.plt <- plot_grid(plimpute, plimpute2, plimpute3, plimpute4, align="v", ncol=1)

#ggsave(FIGURE.5A, plot = impute.plt, path=OUT.PATH, device = pdf, width = 3, height = 6, units = "in")
print("Constructing Figure 5B")
fig.5b.data <- dplyr::filter(STABLE5.DATA, Figure == "Figure_5b")
fig.5b.levels <- c("", "BUSSEMAKER", "ATAC-MOTIF", "YEASTRACT", "NONE-NoTFA")
fig.5b.labels <- c("Gold Standard", "Bussemaker", "ATAC-Motif", "YEASTRACT", "No Priors / TFA")

prior.plt <- ggplot(fig.5b.data, aes(x=Count, y=AUPR, group=Priors)) +
  stat_summary(aes(y=AUPR, group=Priors, linetype=Priors), fun.y=median, geom='line') +
  add_plot_theme() +
  labs(x = "Number of Cells", y = "Median AUPR") +
  scale_x_log10(breaks = c(1, 10, 100, 1000, 10000, 40000)) +
  scale_y_continuous(limits = c(0, NA)) +
  scale_linetype_discrete(limits=fig.5b.levels, labels=fig.5b.labels)

#ggsave(FIGURE.5B, plot = prior.plt, path=OUT.PATH, device = pdf, width = 4, height = 6, dpi = 300, units = "in")
print("Constructing Figure 5C")
dropout.data <- dplyr::filter(STABLE5.DATA, Figure == "Figure_5c")
dropout.levels <- c("ALL", "CSTARVE", "NLIMUREA", "NLIMNH4", "NLIMPRO", "NLIMGLN", "MMETOH", "MMD", "YPETOH", "RAPA", "DIAUXY", "YPD")
dropout.plt <- make_dotplot(plot.data=dropout.data, x.var='Conditions', y.var='AUPR', x.lab="Included Condition", y.lab='AUPR', plot.title="Single Conditions", y.lim=c(0,max(dropout.data$AUPR)), x.axis.log = FALSE) +
  stat_summary(fun.data = mean_sdl, fun.args = list(mult = 1), color = "black",  position = position_nudge(x = 0.25, y = 0)) +
  scale_x_discrete(limits=dropout.levels, labels=c("ALL", CONDITION.LABELS))
#ggsave(FIGURE.5C, plot = dropout.plt, path=OUT.PATH, device = pdf, width = 4, height = 6, dpi = 300, units = "in")
print("Constructing Figure 5D")
mtl.data <- dplyr::filter(STABLE5.DATA, Figure == "Figure_5d")
mtl.data <- dplyr::filter(mtl.data, Priors == "YEASTRACT")
mtl.levels <- c("STL", "MTL")

mtl.aupr.plt <- make_dotplot(mtl.data, 'Tasking', 'AUPR', NULL, 'AUPR', y.lim=c(0,max(mtl.data$AUPR, na.rm = TRUE)), x.axis.log = FALSE) +
  scale_x_discrete(limits=mtl.levels) +
  stat_summary(fun.data = mean_sdl, fun.args = list(mult = 1), color = "black",  position = position_nudge(x = 0.25, y = 0)) +
  theme(axis.text.x = element_blank())

mtl.prec.plt <- make_dotplot(mtl.data, 'Tasking', 'N_PREC', NULL, '# Edges\n(Prec. > 0.5)', y.lim=c(0,max(mtl.data$N_PREC, na.rm = TRUE)), x.axis.log = FALSE) +
  scale_x_discrete(limits=mtl.levels) +
  stat_summary(fun.data = mean_sdl, fun.args = list(mult = 1), color = "black",  position = position_nudge(x = 0.25, y = 0))

figure.5d.plt <- plot_grid(mtl.aupr.plt, mtl.prec.plt, align="v", ncol=1)

figure.5.plt <- plot_grid(impute.plt, prior.plt, dropout.plt, figure.5d.plt, labels = c("A)", "B)", "C)", "D)"), rel_widths = c(0.75, 0.75, 1, 0.75), ncol=4)
ggsave(FIGURE.5.FILE, plot=figure.5.plt, path=OUT.PATH, device = cairo_pdf, width = 8.5, height = 5, units = "in", dpi = FIGURE.DPI)
```
