## Supplementary material for "Gene regulatory network reconstruction using single-cell RNA sequencing of barcoded genotypes in diverse environments": Jackson_2019_Supplemental_Data_1.tar.gz: figure_6.R

```
# Condition-specific networks
if (!exists('SUBNETWORK.FILES')) {
  SUBNETWORK.FILES <- list("CSTARVE"="CSTARVE_signed_network.tsv",
                         "DIAUXY"="DIAUXY_signed_network.tsv",
                         "MMD"="MMD_signed_network.tsv",
                         "MMEtOH"="MMEtOH_signed_network.tsv",
                         "NLIM-GLN"="NLIM-GLN_signed_network.tsv",
                         "NLIM-NH4"="NLIM-NH4_signed_network.tsv",
                         "NLIM-PRO"="NLIM-PRO_signed_network.tsv",
                         "NLIM-UREA"="NLIM-UREA_signed_network.tsv",
                         "RAPA"="RAPA_signed_network.tsv",
                         "YPD"="YPD_signed_network.tsv",
                         "YPEtOH"="YPEtOH_signed_network.tsv")
}

# Output paths
FIGURE.6.FILE <- 'figure_6.pdf'
SUPPLEMENTAL.FIGURE.6.FILE <- 'supplemental_figure_6.pdf'

MAX.TF.NUMBER <- 90
COLUMN.COLORS <- c("red", "gray", "black")
supp.fig.6d.genes <- c("YER062C", "YNL142W", "YHL016C", "YHR070W")

#Number of terms for 6C
if (!exists('NUM.GEO.TERMS')) {NUM.GEO.TERMS <<- 25}
if (!exists('count_data')) {source('figure_common_functions.R')}
if (!exists('sce.data')) {sce.data <<- dimred_for_umap(load_data_into_sce(count.data = dplyr::select(count_data, -META.DATA.COLUMNS), 
                                                                          meta.data = meta_data), 
                                                       random_state=RANDOM.SEED)}

print("Creating Figure 6C-E")
# Read in the learned network
network.learned <- read.table(file.path(WORKING.PATH, NETWORK.PATH, NETWORK.FILE), sep = "\t", header = TRUE, row.names = 1)
# Read in the priors
network.priors <- read.table(file.path(WORKING.PATH, PRIORS.PATH, PRIORS.FILE), sep = "\t", header = TRUE, row.names = 1)

network.learned <- network.learned[rownames(network.learned) %in% rownames(network.priors),]
network.priors <- network.priors[rowSums(network.priors) > 0, ]

# Read in GO terms
go.slim <- read.table(file.path(WORKING.PATH, DATA.PATH, GO.SLIM.FILE), sep="\t")
colnames(go.slim) <- c("ORF", "Gene", "SGDID", "GO_Aspect", "GO_Slim", "GO_ID", "Feature_Type")
go.slim$ORF <- as.character(go.slim$ORF)

# Restrict GO terms to just the terms that we're interested in
go.slim <- go.slim[go.slim$GO_Aspect == "P",]

count.go.from.network <- function(go, learned.data, prior.data, go.axis=0) {

  # Summarize on this axis
  network.summary <- parse.network.and.priors(learned.data, prior.data, parse.axis=go.axis)
  
  # Combine the gene learned/hit/miss counts with the GO Slim data and sum up the results for each go term
  go %>%
    dplyr::left_join(network.summary$learned, by = c("ORF" = "Gene")) %>%
    dplyr::left_join(network.summary$prior.hit, by = c("ORF" = "Gene")) %>%
    dplyr::left_join(network.summary$prior.miss, by = c("ORF" = "Gene")) %>%
    dplyr::mutate(learned = dplyr::coalesce(learned, 0), prior.hit = dplyr::coalesce(prior.hit, 0),
                  prior.miss = dplyr::coalesce(prior.miss, 0)) %>%
    dplyr::mutate(total = learned + prior.hit + prior.miss) %>%
    dplyr::group_by(GO_Slim) %>%
    dplyr::summarise(genes = dplyr::n(), total = sum(total), prior.hit = sum(prior.hit), prior.miss=sum(prior.miss), learned = sum(learned)) -> go.counts
  
  # Get the ordering that we want
  go.plt.order <- order((go.counts$prior.hit + go.counts$learned), decreasing = TRUE)
  go.plt.limit <- as.character(go.counts$GO_Slim[go.plt.order])
  
  # Melt the data for ggplot
  dplyr::select(go.counts, c("GO_Slim", "learned", "prior.hit", "prior.miss")) %>%
    reshape2::melt(id.vars="GO_Slim", variable.name = "model.result", value.name = "edges") -> go.counts
  
  return(list(data=go.counts, order=go.plt.limit))
}

plt.go.data <- function(go.data, drop.go.terms) {
  go.data$data <- dplyr::filter(go.data$data, !GO_Slim %in% drop.go.terms)
  go.data$order <- go.data$order[!go.data$order %in% drop.go.terms]
  
  go.plt <- ggplot(go.data$data, aes(x = GO_Slim, y=edges)) +
    geom_col(aes(fill=model.result)) +
    theme_classic() + 
    theme(axis.text.x = element_text(angle=90, vjust=0.5, hjust=1), 
          plot.title = element_text(face = "bold", hjust=0.5),
          legend.position="none") +
    scale_x_discrete(limits = go.data$order[1:NUM.GEO.TERMS]) +
    scale_fill_manual(values=COLUMN.COLORS, limits=c("learned","prior.hit","prior.miss"), labels = c("Learned", "Recovered", "Lost"), name="Outcome") +
    labs(x="GO Slim Terms", y = "Interactions")
  
  return(go.plt)
}

target.go.data <- count.go.from.network(go.slim, network.learned, network.priors, go.axis=0)
bad.target.go.terms <- c("biological_process", "not_yet_annotated", "other")
target.go.plt <- plt.go.data(target.go.data, bad.target.go.terms) + labs(title="Targets")

regulator.go.data <- count.go.from.network(go.slim, network.learned, network.priors, go.axis=1)
bad.regulator.go.terms <- c("biological_process", "not_yet_annotated", "other", "transcription from RNA polymerase II promoter")
regulator.go.plt <- plt.go.data(regulator.go.data, bad.regulator.go.terms) + labs(title="Regulators")

tf.network.summary <- parse.network.and.priors(network.learned, network.priors, parse.axis = 1)
gene.network.summary <- parse.network.and.priors(network.learned, network.priors)

base::data.frame('Gene' = base::union(colnames(network.learned), colnames(network.priors))) %>%
  dplyr::left_join(tf.network.summary$learned, by = "Gene") %>%
  dplyr::left_join(tf.network.summary$prior.hit, by = "Gene") %>%
  dplyr::left_join(tf.network.summary$prior.miss, by = "Gene") %>%
  dplyr::mutate(learned = dplyr::coalesce(learned, 0), prior.hit = dplyr::coalesce(prior.hit, 0),
                prior.miss = dplyr::coalesce(prior.miss, 0)) %>%
  reshape2::melt(id.vars="Gene", variable.name = "model.result", value.name = "edges") -> tf.counts

tf.plt.order <- tf.network.summary$learned$Gene[order(tf.network.summary$learned$learned, decreasing = TRUE)]
tf.plt.number <- min(sum(tf.network.summary$learned$learned > 0), MAX.TF.NUMBER)

figure.6c.plt <- ggplot(tf.counts, aes(x = Gene, y = edges)) +
  geom_col(aes(fill=model.result)) +
  theme_classic() + 
  theme(axis.text.x = element_text(angle=90, vjust=0.5, hjust=1, size = 8, color = "black", face = "italic"),
        axis.title.y = element_blank(),
        plot.title = element_text(face = "bold", hjust=0.5),
        legend.position="none") +
  scale_x_discrete(limits = tf.plt.order[1:tf.plt.number], labels=yeast.systemic.to.common(tf.plt.order[1:tf.plt.number])) +
  scale_fill_manual(values=COLUMN.COLORS, limits=c("learned","prior.hit","prior.miss"), labels = c("Learned", "Recovered", "Lost"), name="Outcome") +
  labs(x="TFs", y = "Interactions")

figure.6.plt <- plot_grid(figure.6c.plt, plot_grid(target.go.plt, regulator.go.plt, ncol=2, labels = c("i", "ii")), ncol=1, labels=c("C)", "D)"), rel_heights = c(1, 2))
ggsave(FIGURE.6.FILE, plot = figure.6.plt, path=OUT.PATH, device=cairo_pdf, height = 10, width=8.5, units = "in", dpi = FIGURE.DPI)

print("Creating Supplemental Figure 6A")
yeast.gene.names %>%
  dplyr::left_join(gene.network.summary$learned, by = c("SystematicName" = "Gene")) %>%
  dplyr::left_join(gene.network.summary$prior.hit, by = c("SystematicName" = "Gene")) %>%
  dplyr::left_join(gene.network.summary$prior.miss, by = c("SystematicName" = "Gene")) %>%
  dplyr::mutate(learned = dplyr::coalesce(learned, 0), prior.hit = dplyr::coalesce(prior.hit, 0),
                prior.miss = dplyr::coalesce(prior.miss, 0)) %>%
  dplyr::mutate(total = learned + prior.hit + prior.miss, new.total = learned + prior.hit, prior.total = prior.hit + prior.miss) %>%
  base::as.data.frame() -> all.orf.network

as.data.frame(table(all.orf.network$new.total), responseName="New") %>%
  dplyr::left_join(as.data.frame(table(all.orf.network$prior.total), responseName="Prior"), by="Var1") %>%
  dplyr::mutate(Var1 = as.integer(Var1)) -> supp.fig.6a.data

supp.fig.6a.data$New[is.na(supp.fig.6a.data$New)] = 0
supp.fig.6a.data$Prior[is.na(supp.fig.6a.data$Prior)] = 0

supp.data.6a.sums <- colSums(supp.fig.6a.data[supp.fig.6a.data$Var1 >19, ])
supp.fig.6a.data <- supp.fig.6a.data[supp.fig.6a.data$Var1 <= 20,]
supp.fig.6a.data[supp.fig.6a.data$Var1 == 20, 'New'] = supp.data.6a.sums['New']
supp.fig.6a.data[supp.fig.6a.data$Var1 == 20, 'Prior'] = supp.data.6a.sums['Prior']

supp.fig.6a1.hist <- ggplot(supp.fig.6a.data, aes(x=Var1, y=New)) +
  geom_col() +
  scale_x_discrete(limits = c(0:20), breaks = c(0:20), labels = c(0:19, ">20")) +
  scale_y_continuous(limits = c(0,3000)) +
  coord_cartesian(xlim=c(0,20)) +
  labs(x="# Regulators", y="# Genes")

supp.fig.6a2.hist <- ggplot(supp.fig.6a.data, aes(x=Var1, y=Prior)) +
  geom_col() +
  scale_x_discrete(limits = c(0:20), breaks = c(0:20), labels = c(0:19, ">20")) +
  scale_y_continuous(limits = c(0,3000)) +
  coord_cartesian(xlim=c(0,20)) +
  labs(x="# Regulators", y="# Genes")

supp.fig.6a.plt <- plot_grid(supp.fig.6a1.hist, supp.fig.6a2.hist, ncol=1, labels = c("i", "ii"))

print("Creating Supplemental Figure 6B")
supp.fig.6b.data <- as.data.frame(base::colSums(dplyr::select(count_data, -META.DATA.COLUMNS)))
colnames(supp.fig.6b.data) <- "total.reads"
supp.fig.6b.data$SystematicName <- rownames(supp.fig.6b.data)
supp.fig.6b.data$mean.reads <- supp.fig.6b.data$total.reads / base::nrow(count_data)
supp.fig.6b.data <- dplyr::left_join(all.orf.network, supp.fig.6b.data, by = "SystematicName")
supp.fig.6b.corr <- stats::cor(supp.fig.6b.data$mean.reads, supp.fig.6b.data$new.total, method="spearman", use="pairwise.complete.obs")
supp.fig.6b.plt <- ggplot(supp.fig.6b.data, aes(x=rank(mean.reads, ties.method = "first"), y=new.total)) +
  geom_point(size=0.25) +
  geom_smooth(method = "lm", se=FALSE, color="red", linetype="longdash", formula = y~x-1) +
  labs(x="rank(Mean Gene Expression [UMI Count])", y="Learned # TFs Regulating)") +
  geom_label(x=5000, y=13, label=paste0("r2 = ", round(supp.fig.6b.corr, digits=3)), fontface="bold") +
  theme(axis.text.x = element_blank())

print("Creating Supplemental Figure 6C")
subnetworks <- lapply(SUBNETWORK.FILES, function(x) {read.table(file.path(NETWORK.PATH, x), sep = "\t", header = TRUE, row.names = 1)})

activator.learned <- base::data.frame(base::matrix(0, nrow = base::nrow(network.priors), ncol = base::ncol(network.priors)))
rownames(activator.learned) <- rownames(network.priors)
colnames(activator.learned) <- colnames(network.priors)

repressor.learned <- base::data.frame(base::matrix(0, nrow = base::nrow(network.priors), ncol = base::ncol(network.priors)))
rownames(repressor.learned) <- rownames(network.priors)
colnames(repressor.learned) <- colnames(network.priors)

for (cond in names(subnetworks)) {
  subnet <- subnetworks[[cond]]
  roverlap <- rownames(network.priors)[rownames(network.priors) %in% rownames(subnet)]
  coverlap <- colnames(network.priors)[colnames(network.priors) %in% colnames(subnet)]
  activator.learned[roverlap, coverlap] <- activator.learned[roverlap, coverlap] + (subnet > 0)[roverlap, coverlap]
  repressor.learned[roverlap, coverlap] <- repressor.learned[roverlap, coverlap] + (subnet < 0)[roverlap, coverlap]
}

network.learned <- network.learned[rownames(network.learned) %in% rownames(network.priors),]
network.learned[network.learned == 0] <- NA
network.learned <-reshape2::melt(as.matrix(network.learned), na.rm = TRUE)
colnames(network.learned) <- c("target", "regulator", "int")

activator.learned[activator.learned == 0] <- NA
repressor.learned[repressor.learned == 0] <- NA
network.data <- list(activator = reshape2::melt(as.matrix(activator.learned), na.rm = TRUE),
                     repressor = reshape2::melt(as.matrix(repressor.learned), na.rm = TRUE))
colnames(network.data[["activator"]]) <- c("target", "regulator", "act.int")
colnames(network.data[["repressor"]]) <- c("target", "regulator", "rep.int")

network.learned %>%
  dplyr::left_join(network.data[["activator"]], by=c("target", "regulator")) %>%
  dplyr::left_join(network.data[["repressor"]], by=c("target", "regulator")) %>%
  dplyr::mutate(act.int = dplyr::coalesce(act.int, 0), rep.int = dplyr::coalesce(rep.int, 0)) -> network.data

network.data %>%
  dplyr::filter(int == 1) %>%
  dplyr::group_by(regulator, act.int) %>%
  dplyr::summarise(num=dplyr::n()) %>%
  base::as.data.frame() -> activation.data

network.data %>%
  dplyr::filter(int == -1) %>%
  dplyr::group_by(regulator, rep.int) %>%
  dplyr::summarise(num=dplyr::n()) %>%
  base::as.data.frame() -> repression.data

network.data %>%
  dplyr::group_by(regulator) %>%
  dplyr::summarise(num=dplyr::n()) -> tf.int.count

tf.plt.order <- tf.int.count$regulator[order(tf.int.count$num, decreasing = TRUE)]

supp.fig.6c.i.plt <- ggplot(activation.data, aes(x=regulator, y=num)) +
  geom_col(aes(fill=act.int)) +
  theme_classic() + 
  theme(axis.text.x = element_text(angle=90, vjust=0.5, hjust=1, size = 8, color = "black", face = "italic"),
        axis.title.y = element_blank(),
        plot.title = element_text(face = "bold", hjust=0.5)) +
  scale_x_discrete(limits = tf.plt.order[1:90], 
                   labels=yeast.systemic.to.common(tf.plt.order[1:90])) +
  scale_y_continuous(limit=c(0, 1700)) +
  scale_fill_viridis() +
  labs(x="TFs", y = "Interactions")

supp.fig.6c.ii.plt <- ggplot(repression.data, aes(x=regulator, y=num)) +
  geom_col(aes(fill=rep.int)) +
  theme_classic() + 
  theme(axis.text.x = element_text(angle=90, vjust=0.5, hjust=1, size = 8, color = "black", face = "italic"),
        axis.title.y = element_blank(),
        plot.title = element_text(face = "bold", hjust=0.5)) +
  scale_x_discrete(limits = tf.plt.order[1:90], 
                   labels=yeast.systemic.to.common(tf.plt.order[1:90])) +
  scale_y_continuous(limit=c(0, 1700)) +
  scale_fill_viridis() +
  labs(x="TFs", y = "Interactions")

print("Creating Supplemental Figure 6D")
supp.fig.6d.conditions <- c("YPDRapa")
supp.fig.6d.plt <- plt.gene.violins(sce.data, supp.fig.6d.genes, conditions = supp.fig.6d.conditions) +
  facet_wrap(~gene, labeller = as_labeller(yeast.systemic.to.common), ncol = 4) +
  labs(x="Genotype", y="log2(Expression)")

supp.fig.6 <- plot_grid(plot_grid(supp.fig.6a.plt, supp.fig.6b.plt, ncol=2, labels = c("A)", "B)")), 
                        supp.fig.6c.i.plt, supp.fig.6c.ii.plt, supp.fig.6d.plt, 
                        ncol=1, labels = c("", "C)", "", "D)"), rel_heights = c(1, 0.75, 0.75, 1))
ggsave(SUPPLEMENTAL.FIGURE.6.FILE, plot = supp.fig.6, path=OUT.PATH, device=cairo_pdf, height = 9, width=8.5, units = "in", dpi = FIGURE.DPI)
```
