## Supplementary material for "Gene regulatory network reconstruction using single-cell RNA sequencing of barcoded genotypes in diverse environments": Jackson_2019_Supplemental_Data_1.tar.gz: figure_7.R

```
FIG.7A.CONDITIONS <- c("YPD", "YPDRapa", "AmmoniumSulfate", "CStarve")
FIG.7A.TFs <- c("YFL021W", "YJL110C", "YER040W", "YKR034W", "YIR023W", "YEL009C")
FIG.7A.TF.MAP <- structure(c("gat1", "gzf3", "gln3", "dal80", "dal81", "gcn4"), names=FIG.7A.TFs)

FIG.7B.TFs <- c("YOR372C", "YIL131C", "YNL068C", "YMR043W", "YER111C", "YDR146C")
SUPP.FIG.7.CC.TARGETS <- c("YKL164C", "YER124C", "YHR143W", "YDR224C", "YBR009C", "YNL300W")

FIGURE.7A.FILE <- 'figure_7a.pdf'
FIGURE.7B.FILE <- 'figure_7b.png'
SUPP.FIG.7.FILE <- 'supplemental_figure_7.png'

range.norm <- function(x, y) {(x - min(y))/(max(y) - min(y))}

if (!exists('count_data')) {source('figure_common_functions.R')}
if (!exists('sce.data')) {sce.data <<- dimred_for_umap(load_data_into_sce(count.data = dplyr::select(count_data, -META.DATA.COLUMNS), 
                                                                          meta.data = meta_data), 
                                                       random_state=RANDOM.SEED)}

print("Constructing Figure 7A")
network.learned <- read.table(file.path(WORKING.PATH, NETWORK.PATH, NETWORK.FILE), sep = "\t", header = TRUE, row.names = 1)
network.priors <- read.table(file.path(WORKING.PATH, PRIORS.PATH, PRIORS.FILE), sep = "\t", header = TRUE, row.names = 1)
count.for.activity <- as.matrix(t(dplyr::select(count_data, -META.DATA.COLUMNS)[, rownames(network.learned)]))
activities <- t(tfa(as.matrix(network.learned), count.for.activity))
SingleCellExperiment::reducedDim(sce.data, "activity") <- activities
SingleCellExperiment::reducedDim(sce.data, "scaledactivity") <- scale(activities)

fig.7a.plt.list <- list()
for (tf.name in FIG.7A.TFs) {
  base::as.data.frame(activities)[, tf.name, drop=FALSE] %>%
    stats::setNames("TF") %>%
    base::cbind(meta_data) %>%
    dplyr::filter(Genotype_Group != FIG.7A.TF.MAP[[tf.name]]) %>%
    dplyr::select(-c(tenXBarcode, Genotype_Group, Genotype, Replicate)) %>%
    dplyr::mutate(TF = scale(TF)) %>%
    dplyr::group_by(Condition) %>%
    dplyr::summarise_all(mean) %>%
    dplyr::mutate(TF = base::round(TF, digits=4)) -> activity.by.condition
  
  fig.7a.plt.list[[tf.name]] <- ggplot(activity.by.condition, aes(x=Condition, y=TF)) +
    geom_col(fill = "black") +
    theme_classic() +
    scale_x_discrete(limits = rev(FIG.7A.CONDITIONS), 
                     breaks = rev(FIG.7A.CONDITIONS), 
                     labels=rev(translate.condition.name(FIG.7A.CONDITIONS))) +
    scale_y_continuous(limit=c(-1, 1), breaks = c(-1, 0, 1), labels = c(-1, 0, 1)) +
    labs(x = paste0(yeast.systemic.to.common(tf.name), " Activity"), y = "Condition") +
    theme(axis.text = element_text(size=10)) +
    coord_flip()
}
fig.7a.plt <- plot_grid(plotlist = fig.7a.plt.list, ncol=3)
ggsave(FIGURE.7A.FILE, plot = fig.7a.plt, path=OUT.PATH, device=cairo_pdf, height = 4, width=6, units = "in", dpi = FIGURE.DPI)

print("Constructing Figure 7B")
sce.ypd <- sce.data[, sce.data$Condition == "YPD"]
sce.ypd <- normalizeSCE(sce.ypd)
sce.ypd <- runPCA(sce.ypd, ncomponents=50, ntop=Inf, method='irlba', exprs_values='logcounts')
sce.ypd <- runUMAP(sce.ypd, random_state=RANDOM.SEED, min_dist=0.1, use_dimred='PCA')

fig7b.plt.lst <- lapply(FIG.7B.TFs, function(x) {umap.plot(sce.ypd, gene = x, exp.name="scaledactivity", exp.type="reduceddim", alpha=0.5, size=0.05, color.title='Condition') + 
  scale_colour_viridis(option='B', limits = c(-4,4), oob = squish) +
  theme(legend.position = "none", axis.title = element_blank(), axis.text = element_blank(),
        plot.title = element_text(size=12, face = "bold.italic", hjust = 0.5)) +
  labs(title = yeast.systemic.to.common(x))})

fig7b.plt <- plot_grid(plotlist=fig7b.plt.lst, ncol=6)
ggsave(FIGURE.7B.FILE, plot = fig7b.plt, path=OUT.PATH, device="png", height = 2, width=8.5, units = "in", dpi = FIGURE.DPI)

print("Constructing Supplemental Figure 7")
supp.fig.7a.plt.list <- lapply(FIG.7A.TFs, function(x) {umap.plot(sce.ypd, gene = x, exp.name="scaledactivity", exp.type="reduceddim", alpha=0.5, size=0.05, color.title='Condition') + 
    scale_colour_viridis(option='B', limits = c(-4,4), oob = squish) +
    theme(legend.position = "none", axis.title = element_blank(), axis.text = element_blank(),
          plot.title = element_text(size=12, face = "bold.italic", hjust = 0.5)) +
    labs(title = yeast.systemic.to.common(x))})

supp.fig.7b.plt.list <- lapply(FIG.7B.TFs, function(x) {umap.plot(sce.ypd, gene = x, exp.name="logcounts", alpha=0.5, size=0.05, color.title='Condition') + 
    scale_colour_viridis(option='B', limits = c(0,2), oob = squish) +
    theme(legend.position = "none", axis.title = element_blank(), axis.text = element_blank(),
          plot.title = element_text(size=12, face = "bold.italic", hjust = 0.5)) +
    labs(title = yeast.systemic.to.common(x))})

supp.fig.7c.plt.list <- lapply(SUPP.FIG.7.CC.TARGETS, function(x) {umap.plot(sce.ypd, gene = x, exp.name="logcounts", alpha=0.5, size=0.05, color.title='Condition') + 
    scale_colour_viridis(option='B', limits = c(0,2), oob = squish) +
    theme(legend.position = "none", axis.title = element_blank(), axis.text = element_blank(),
          plot.title = element_text(size=12, face = "bold.italic", hjust = 0.5)) +
    labs(title = yeast.systemic.to.common(x))})

supp.fig.7a.plt <- plot_grid(plotlist=supp.fig.7a.plt.list, ncol=6)
supp.fig.7b.plt <- plot_grid(plotlist=supp.fig.7b.plt.list, ncol=6)
supp.fig.7c.plt <- plot_grid(plotlist=supp.fig.7c.plt.list, ncol=6)

supp.fig.7 <- plot_grid(supp.fig.7a.plt, fig7b.plt, supp.fig.7b.plt, supp.fig.7c.plt, ncol=1, labels=c("A)", "B)", "C)"))
ggsave(SUPP.FIG.7.FILE, plot = supp.fig.7, path=OUT.PATH, device="png", height = 7, width=8.5, units = "in", dpi = FIGURE.DPI)
```
