## Supplementary material for "Gene regulatory network reconstruction using single-cell RNA sequencing of barcoded genotypes in diverse environments": Jackson_2019_Supplemental_Data_1.tar.gz: figure_common_functions.R

```
# Bioconductor
require(scater)
require(scran)
require(SingleCellExperiment)
require(SummarizedExperiment)
require(BiocParallel)

# Tidyverse
require(data.table)
require(stringr)
require(dplyr)
require(reshape2)

# Plotting
require(igraph)
require(ggplot2)
require(ggridges)
require(scales)
require(viridis)
require(RColorBrewer)
require(cowplot)

# Math
require(umap)
require(corpcor)

# Set a bunch of defaults if they're not loaded by the main script (so each figure script can be fun separately)

# File paths (set if they are not already in the global space)
if(!exists('WORKING.PATH')) {WORKING.PATH <<- "."}
if(!exists('COUNTS.FILE')) {COUNTS.FILE <<- "103118_SS_Data.tsv.gz"}
if(!exists('GENE.NAME.FILE')) {GENE.NAME.FILE <<- 'yeast_gene_names.tsv'}
if(!exists('NETWORK.FILE')) {NETWORK.FILE <<- 'signed_network.tsv'}
if(!exists('PRIORS.FILE')) {PRIORS.FILE <<- 'YEASTRACT_priors_20181118.tsv.gz'}
if(!exists('GO.SLIM.FILE')) {GO.SLIM.FILE <<- 'go_slim_mapping.tab'}
if(!exists('OUT.PATH')) {OUT.PATH <<- 'figures'}
if(!exists('DATA.PATH')) {DATA.PATH <<- 'data'}
if(!exists('PRIORS.PATH')) {PRIORS.PATH <<- 'priors'}
if(!exists('NETWORK.PATH')) {NETWORK.PATH <<- 'network'}
if(!exists('RANDOM.SEED')) {RANDOM.SEED <<- 42}
if(!exists('MP.CORES')) {MP.CORES <<- 15}
if(!exists('FIGURE.DPI')) {FIGURE.DPI <<- 300}
if(!exists('META.DATA.COLUMNS')) {META.DATA.COLUMNS <<- c("Condition", "Genotype", "Genotype_Group", "tenXBarcode", "Replicate")}
if (!exists('FIG4.P.ADJ.THRESHOLD')) {FIG4.P.ADJ.THRESHOLD <<- 0.05}
if (!exists('FIG4.LFC.THRESHOLD')) {FIG4.LFC.THRESHOLD <<- log2(1.5)}

# Set levels, labels, and colors for categorical variables CONDITION and GENOTYPE
if(!exists('CONDITION.COLORS')) {CONDITION.COLORS <<- c('lightseagreen', 'darkgoldenrod1', 'darkolivegreen2', 'deepskyblue2', 'darkorchid2', 'deeppink1', 'forestgreen', 'green2', 'red4', 'palevioletred1', 'slategray3')}
if(!exists('CONDITION.LEVELS')) {CONDITION.LEVELS <<- c("CStarve", "Urea", "AmmoniumSulfate", "Proline", "Glutamine", "MinimalEtOH", "MinimalGlucose", "YPEtOH", "YPDRapa", "YPDDiauxic", "YPD")}
if(!exists('CONDITION.LABELS')) {CONDITION.LABELS <<- c("CSTARVE", "NLIM-UREA", "NLIM-NH4", "NLIM-PRO", "NLIM-GLN", "MMEtOH", "MMD", "YPEtOH", "RAPA", "DIAUXY", "YPD")}
if(!exists('GENOTYPE.COLORS')) {GENOTYPE.COLORS <<- brewer.pal(12, 'Set3')}
if(!exists('GENOTYPE.LEVELS')) {GENOTYPE.LEVELS <<- c("stp2", "stp1", "rtg3", "rtg1", "gcn4",  "dal82", "dal81", "dal80", "gat1", "gln3", "gzf3", "WT(ho)")}
if(!exists('GENOTYPE.LABELS')) {GENOTYPE.LABELS <<- c("Δstp2", "Δstp1", "Δrtg3", "Δrtg1", "Δgcn4",  "Δdal82", "Δdal81", "Δdal80", "Δgat1", "Δgln3", "Δgzf3", "WT")}

# Set WD and create output path
setwd(WORKING.PATH)
dir.create(OUT.PATH, showWarnings = FALSE)

# Read in yeast gene names and create a translator to turn a vector of systemic names into a vector of common names
# For labelling purposes

yeast.gene.names <- read.table(file.path(WORKING.PATH, DATA.PATH, GENE.NAME.FILE), sep = "\t", quote="", header=TRUE, stringsAsFactors = FALSE)
yeast.gene.names[yeast.gene.names == ""] = NA

yeast.systemic.to.common <- function(name.vec) {
  translated.names <- match(name.vec, yeast.gene.names$SystematicName, nomatch=NA)
  translated.names <- yeast.gene.names[translated.names, "Name"]
  no.translation <- is.na(translated.names)
  translated.names[no.translation] <- name.vec[no.translation]
  return(translated.names)
}

translate.condition.name <- function(cond) {structure(CONDITION.LABELS, names=CONDITION.LEVELS)[cond]}

fix_imported_data_frame <- function(broken.data.frame) {
  rownames(broken.data.frame) <- gsub(".", "-", rownames(broken.data.frame), fixed = TRUE)
  colnames(broken.data.frame) <- gsub(".", "-", colnames(broken.data.frame), fixed = TRUE)
  rownames(broken.data.frame)[which(rownames(broken.data.frame) == "X15S_rRNA")] =  "15S_rRNA"
  rownames(broken.data.frame)[which(rownames(broken.data.frame) == "X21S_rRNA")] =  "21S_rRNA"
  return(broken.data.frame)
}

# The following functions manipulate the SingleCellExperiment data container for SCRAN/SCATER

load_data_into_sce <- function(count.data, meta.data, genes.are.columns=TRUE) {
  
  if (genes.are.columns) {count.data <- t(count.data)}

  sce <- SingleCellExperiment::SingleCellExperiment(list(counts=count.data))
  sce$Genotype_Replicate = meta.data$Genotype
  sce$Genotype_Group = meta.data$Genotype_Group
  sce$Condition = meta.data$Condition
  
  return(sce)
}

dimred_for_umap <- function(sce, batch="Condition", do.multibatchnorm=TRUE, random_state=42) {
  if(do.multibatchnorm) {sce <- apply.multibatchnorm(sce, batch = batch)}
  else {SummarizedExperiment::assay(sce, "logcounts") <- SummarizedExperiment::assay(sce, "counts")}
  sce <- apply.mpca(sce, batch = batch)
  print("Reducing dimensionality with UMAP")
  umap.data <- as.matrix(SingleCellExperiment::reducedDim(sce, 'MPCA'))
  umap.result <- umap::umap(umap.data, method = "umap-learn", random_state = random_state)
  SingleCellExperiment::reducedDim(sce, 'UMAP') <- umap.result$layout
  print("Processing Complete")
  return(sce)
}

apply.multibatchnorm <- function(sce, batch = "Condition") {
  print("Normalizing batches with multibatchNorm...")
  rescaled <- scran::multiBatchNorm(sce[, sce[[batch]] == 'Glutamine'], sce[, sce[[batch]] == 'Proline'], 
                                     sce[, sce[[batch]] == 'AmmoniumSulfate'], sce[, sce[[batch]] == 'Urea'], 
                                     sce[, sce[[batch]] == 'YPD'], sce[, sce[[batch]] == 'YPDRapa'],
                                     sce[, sce[[batch]] == 'CStarve'], sce[, sce[[batch]] == 'MinimalGlucose'], 
                                     sce[, sce[[batch]] == 'MinimalEtOH'], sce[, sce[[batch]] == 'YPDDiauxic'], 
                                     sce[, sce[[batch]] == 'YPEtOH'])
  
  SummarizedExperiment::assay(sce, "logcounts") <- do.call('cbind', 
                                                           lapply(rescaled, 
                                                                  function(x){SummarizedExperiment::assays(x)[['logcounts']]}))
  return(sce)
}

apply.mpca <- function(sce, batch = "Condition") {
  print("Reducing dimensionality with multibatchPCA...")
  multibatch.pca <- scran::multiBatchPCA(sce[, sce[[batch]] == 'Glutamine'], sce[, sce[[batch]] == 'Proline'], 
                                  sce[, sce[[batch]] == 'AmmoniumSulfate'], sce[, sce[[batch]] == 'Urea'], 
                                  sce[, sce[[batch]] == 'YPD'], sce[, sce[[batch]] == 'YPDRapa'],
                                  sce[, sce[[batch]] == 'CStarve'], sce[, sce[[batch]] == 'MinimalGlucose'], 
                                  sce[, sce[[batch]] == 'MinimalEtOH'], sce[, sce[[batch]] == 'YPDDiauxic'], 
                                  sce[, sce[[batch]] == 'YPEtOH'],
                                  approximate=TRUE, subset.row=NULL, d=50,
                                  assay.type='logcounts', get.spikes=FALSE, 
                                  BPPARAM = BiocParallel::MulticoreParam(MP.CORES))
  
  multibatch.pca <- do.call('rbind', multibatch.pca)
  reducedDim(sce, 'MPCA') <- multibatch.pca
  return(sce)
}

apply.snn.cluster <- function(sce) {
  snn.gr <- scran::buildSNNGraph(sce, use.dimred='PCA')
  clusters <- igraph::cluster_louvain(snn.gr)
  sce$cluster <- as.factor(clusters$membership)
  return(sce)
}

umap.plot <- function (sce.exp, color.by=NULL, color.title=NULL, gene=NULL, exp.name='logcounts', exp.type="assay", alpha=0.5, size=1, txt.size=10, shuffle.data = TRUE) {
  
  df <- as.data.frame(reducedDim(sce.exp, 'UMAP'))
  colnames(df) <- c('UMAP1', 'UMAP2')
  
  if (!is.null(color.by)) {
    df$color <- sce.exp[[color.by]]
    if (is.null(color.title)) { 
      color.title <- color.by
    }
  }
  
  if (!is.null(gene)) {
    if (exp.type == "assay") {df$color <- as.vector(colSums(SummarizedExperiment::assay(sce.exp[gene, ], exp.name)))}
    if (exp.type == "reduceddim") {df$color <- as.vector(colSums(t(SingleCellExperiment::reducedDim(sce.exp, exp.name)[, gene])))}
    if (is.null(color.title)) { 
      color.title <- gene
    }
  }
  
  if (shuffle.data) {df <- df[sample(nrow(df)),]}
  
  plt <- ggplot(df, aes(x=UMAP1, y=UMAP2)) + 
    geom_point(aes(color=color), alpha=alpha, size=size) + 
    theme_classic() + labs(color=color.title) +
    theme(panel.grid.major=element_blank(), 
          panel.grid.minor=element_blank(),
          text=element_text(size=txt.size)) 
  return(plt)
}

# Plot gene expression distributions as a violin plot from a SCE object
plt.gene.violins <- function(sce.exp, gene.id, conditions=NULL, exp.name='logcounts') {
  plt.data <- extract.expression.data(sce.exp, gene.id, exp.name = exp.name)
  
  if (!is.null(conditions)) {
    plt.data <- plt.data[plt.data$condition %in% conditions, ]
  }
  
  plt.data <- reshape2::melt(plt.data, id.vars = c("condition", "genotype"), variable.name = "gene", value.name = "count")
  
  plt <- ggplot(plt.data, aes(x=genotype, y=count)) + 
    geom_violin(aes(fill=genotype), scale = 'width') +
    theme_classic() + 
    scale_fill_manual(labels = GENOTYPE.LABELS, breaks = GENOTYPE.LEVELS, values = GENOTYPE.COLORS) +
    scale_x_discrete(labels = rev(GENOTYPE.LABELS), limits = rev(GENOTYPE.LEVELS)) +
    geom_boxplot(outlier.shape = NA, width=0.25) +
    theme(axis.text.x = element_text(size = 8, angle=90, vjust=0.5), axis.title.x = element_text(size = 12),
          axis.text.y = element_text(size = 10), axis.title.y = element_text(size = 12),
          legend.text = element_text(size = 10), legend.title = element_text(size = 12, face = 'bold'),
          plot.title = element_text(size = 12, face = "bold", hjust=0.5),
          panel.grid.major=element_blank(), panel.grid.minor=element_blank(),
          strip.background = element_blank(), strip.text = element_text(size=10, face="bold"),
          legend.position = "none")
  return(plt)
}


# Extract gene expression data and cluster identity from a list of SCE objects and return it as a single data.frame
extract.expression.clusters.from.list <- function(list.sce.obj, genes, exp.name="logcounts", include.names=NULL) {
  sce.df <- NULL
  if(is.null(include.names)) {include.names <- names(list.sce.obj)}
  for (cond.name in include.names) {
    df <- extract.expression.clusters(list.sce.obj[[cond.name]], genes, exp.name=exp.name)
    sce.df <- rbind(sce.df, df)
  }
  return(sce.df)
}

# Extract gene expression data and cluster identity from a SCE object and return it as a data.frame
extract.expression.clusters <- function(sce.obj, genes, exp.name="logcounts") {
  sce.df <- extract.expression.data(sce.obj, genes, exp.name=exp.name)
  sce.df$clusters <- sce.obj[['cluster']]
  return(sce.df)
}

# Extract gene expression data from a SCE object and return it as a data.frame
extract.expression.data <- function(sce.obj, genes, exp.name="logcounts") {
  sce.df <- as.data.frame(t(SummarizedExperiment::assay(sce.obj[genes, ], exp.name)))
  sce.df$condition <- sce.obj$Condition
  sce.df$genotype <- sce.obj$Genotype_Group
  return(sce.df)
}

# Extract genotype and cluster data from a SCE object
extract.genotype.count <- function(list.sce.obj, include.names=NULL) {
  sce.df <- NULL
  if(is.null(include.names)) {include.names <- names(list.sce.obj)}
  for (cond.name in include.names) {
    sce.obj <- list.sce.obj[[cond.name]]
    df <- data.frame(cells = colnames(sce.obj))
    df$clusters <- sce.obj$cluster
    df$genotype <- sce.obj$Genotype_Group
    df$condition <- cond.name
    sce.df <- rbind(sce.df, df)
  }
  return(sce.df)
}

# Extract the marker genes which drive clustering from a list of SCE objects and return them as a single data.frame
extract.markers <- function(list.sce.obj, marker.cutoff.num=20, marker.cutoff.FDR=0.1, exp.name='logcounts') {
  sce.df <- NULL
  for (cond.name in names(list.sce.obj)) {
    sce.obj <- list.sce.obj[[cond.name]]
    markers.out <- scran::findMarkers(sce.obj, sce.obj$cluster, direction='up', pval.type='any', assay.type=exp.name, lfc=0.25)
    markers <- NULL
    for (i in 1:length(markers.out)) {
      markers.tmp <- as.data.frame(markers.out[[i]])
      markers.tmp$cluster <- i
      markers.tmp <- markers.tmp[markers.tmp$FDR < marker.cutoff.FDR, c('Top', 'p.value', 'FDR', 'cluster')]
      markers.tmp$Gene <- rownames(markers.tmp)
      markers <- rbind(markers, markers.tmp)
    }
    markers$condition <- cond.name
    markers <- markers[markers$Top < marker.cutoff.num, ]
    markers <- markers[order(markers$FDR), ]
    markers <- markers[!duplicated(markers$Gene), ]
    markers <- markers[order(markers$cluster), ]
    sce.df <- rbind(sce.df, markers)
  }
  return(sce.df)
}

# Summarize a network based on what we've learned compared to priors
parse.network.and.priors <- function(learned.data, prior.data, parse.axis=0) {
  # Shrink the priors to match the network we learned
  shrunk.priors <- prior.data
  shrunk.priors[colnames(learned.data)[!colnames(learned.data) %in% colnames(prior.data)]] <- 0
  shrunk.priors <- shrunk.priors[rownames(learned.data), intersect(colnames(learned.data), colnames(shrunk.priors))]
  shrunk.priors[is.na(shrunk.priors)] <- 0
  
  if (parse.axis == 0) {sum.func <- base::rowSums}
  else if (parse.axis == 1) {sum.func <- base::colSums}
  else {return(FALSE)}
  
  # Figure out what new stuff we predicted
  gene.interaction.counts.learned <- base::as.data.frame(sum.func((learned.data != 0) & (shrunk.priors == 0)))
  colnames(gene.interaction.counts.learned) <- "learned"
  gene.interaction.counts.learned$Gene <- rownames(gene.interaction.counts.learned)
  
  # Figure out which priors we predicted
  gene.interaction.counts.priors.hit <- base::as.data.frame(sum.func((learned.data != 0) & (shrunk.priors != 0)))
  colnames(gene.interaction.counts.priors.hit) <- "prior.hit"
  gene.interaction.counts.priors.hit$Gene <- rownames(gene.interaction.counts.priors.hit)
  
  # Figure out which priors we missed (not off the shrunken priors, off the whole thing)
  expanded.network <- learned.data
  expanded.network[colnames(prior.data)[!colnames(prior.data) %in% colnames(learned.data)]] <- 0
  expanded.network <- expanded.network[rownames(prior.data), colnames(prior.data)]
  expanded.network[is.na(expanded.network)] <- 0
  
  gene.interaction.counts.priors.missed <- base::as.data.frame(sum.func((prior.data != 0 & expanded.network == 0)))
  colnames(gene.interaction.counts.priors.missed) <- "prior.miss"
  gene.interaction.counts.priors.missed$Gene <- rownames(gene.interaction.counts.priors.missed)
  
  return(list(learned=gene.interaction.counts.learned,
              prior.hit=gene.interaction.counts.priors.hit,
              prior.miss=gene.interaction.counts.priors.missed))
}

# Calculate transcription factor activity from a network and an expression matrix
tfa <- function(prior, exp.mat, noself=TRUE, dup.self=TRUE) {
  tfwt <- apply(prior != 0, 2, sum) > 0
  
  duplicates <- c()
  if (dup.self) {
    duplicates <- duplicated(prior[, tfwt], MARGIN=2) |
      duplicated(prior[, tfwt], MARGIN=2, fromLast = TRUE)
    duplicates <- colnames(prior)[tfwt][duplicates]
  }
  
  tfs <- setdiff(colnames(prior), duplicates)
  tfs <- intersect(tfs, rownames(prior))
  if (noself) {
    diag(prior[tfs, tfs]) <- 0
  }
  
  activities <- matrix(0, ncol(prior), ncol(exp.mat), 
                       dimnames=list(colnames(prior), colnames(exp.mat)))
  
  if (any(tfwt)) {
    activities[tfwt, ] <- corpcor::pseudoinverse(prior[, tfwt, drop=FALSE]) %*% exp.mat
  }
  
  use.exp <- intersect(colnames(prior)[!tfwt], rownames(exp.mat))
  activities[use.exp, ] <- exp.mat[use.exp, ]
  
  return(activities)
}

cat("Loading data matrix: ", file.path(WORKING.PATH, DATA.PATH, COUNTS.FILE), "\n")
if (!exists('count_data')) {count_data <<- fix_imported_data_frame(read.table(gzfile(file.path(WORKING.PATH, DATA.PATH, COUNTS.FILE)), header=TRUE, sep="\t"))}
if (!exists('meta_data')) {meta_data <<- count_data[META.DATA.COLUMNS]}
```
