## Supplementary material for "Gene regulatory network reconstruction using single-cell RNA sequencing of barcoded genotypes in diverse environments": Jackson_2019_Supplemental_Data_1.tar.gz: jackson_2019_figures.R

```
# File paths
WORKING.PATH <<- "."
OUT.PATH <<- 'figures'
DATA.PATH <<- 'data'
PRIORS.PATH <<- 'priors'
NETWORK.PATH <<- 'network'

# File names
COUNTS.FILE <<- "103118_SS_Data.tsv.gz"
GENE.NAME.FILE <<- "yeast_gene_names.tsv"
STABLE5.FILE <<- "STable5.tsv"
NETWORK.FILE <<- "signed_network.tsv"
PRIORS.FILE <<- 'YEASTRACT_priors_20181118.tsv.gz'
GO.SLIM.FILE <<- 'go_slim_mapping.tab'

IMPUTED.DATA.FILES <<- list(Unmodified=list("103118_SS_Data.tsv.gz", TRUE),
                            MAGIC=list("MAGIC_DATA.tsv.gz", FALSE),
                            scImpute=list("SCIMPUTE_DATA.tsv.gz", FALSE),
                            VIPER=list("VIPER_DATA.tsv.gz", TRUE))

SUBNETWORK.FILES <<- list("CSTARVE"="CSTARVE_signed_network.tsv",
                         "DIAUXY"="DIAUXY_signed_network.tsv",
                         "MMD"="MMD_signed_network.tsv",
                         "MMEtOH"="MMEtOH_signed_network.tsv",
                         "NLIM-GLN"="NLIM-GLN_signed_network.tsv",
                         "NLIM-NH4"="NLIM-NH4_signed_network.tsv",
                         "NLIM-PRO"="NLIM-PRO_signed_network.tsv",
                         "NLIM-UREA"="NLIM-UREA_signed_network.tsv",
                         "RAPA"="RAPA_signed_network.tsv",
                         "YPD"="YPD_signed_network.tsv",
                         "YPEtOH"="YPEtOH_signed_network.tsv")

# RNG Seed
RANDOM.SEED <<- 42
MP.CORES <<- 15

# Image Settings
FIGURE.DPI <<- 300

# BioConductor single-cell packages
require(scater)
require(scran)
require(SingleCellExperiment)
require(SummarizedExperiment)
require(BiocParallel)

# Tidyverse
require(data.table)
require(dplyr)
require(reshape2)
require(stringr)

# Plotting
require(igraph)
require(ggplot2)
require(ggridges)
require(scales)
require(cowplot)
require(RColorBrewer)
require(viridis)

# Create the output path if needed
setwd(WORKING.PATH)
dir.create(OUT.PATH, showWarnings = FALSE)

# Set common color and labeling schemes
CONDITION.COLORS <<- c('lightseagreen', 'darkgoldenrod1', 'darkolivegreen2', 'deepskyblue2', 'darkorchid2', 'deeppink1', 'forestgreen', 'green2', 'red4', 'palevioletred1', 'slategray3') #, 'navy', 'black'
CONDITION.LEVELS <<- c("CStarve", "Urea", "AmmoniumSulfate", "Proline", "Glutamine", "MinimalEtOH", "MinimalGlucose", "YPEtOH", "YPDRapa", "YPDDiauxic", "YPD")
CONDITION.LABELS <<- c("CSTARVE", "NLIM-UREA", "NLIM-NH4", "NLIM-PRO", "NLIM-GLN", "MMEtOH", "MMD", "YPEtOH", "RAPA", "DIAUXY", "YPD")
GENOTYPE.COLORS <<- brewer.pal(12, 'Set3')
GENOTYPE.LEVELS <<- c("stp2", "stp1", "rtg3", "rtg1", "gcn4",  "dal82", "dal81", "dal80", "gat1", "gln3", "gzf3", "WT(ho)")
GENOTYPE.LABELS <<- c("Δstp2", "Δstp1", "Δrtg3", "Δrtg1", "Δgcn4",  "Δdal82", "Δdal81", "Δdal80", "Δgat1", "Δgln3", "Δgzf3", "WT")

# Define any metadata columns in the data file
META.DATA.COLUMNS <<- c("Condition", "Genotype", "Genotype_Group", "tenXBarcode", "Replicate")

## Load common functions and the count matrix ##
source('figure_common_functions.R')

## Figure specific settings ##

# Set significance thresholds for Figure 4 DESeq
FIG4.P.ADJ.THRESHOLD <<- 0.05
FIG4.LFC.THRESHOLD <<- log2(1.5)

# Number of GO terms to include in 6C
FIG.6C.NUM.TERMS <<- 25

## Run the figure scripts ##

cat("Creating Figure 2 Panels\n")
source('figure_2.R')

cat("Creating Supplemental Figure 2 Panels\n")
source('supplemental_figure_2.R')

cat("Creating Figure 3 & Supplemental Figure 3 Panels\n")
source('figure_3.R')

cat("Creating Figure 4 Panels\n")
source('figure_4.R')

cat("Creating Supplemental Figure 4 Panels\n")
source('supplemental_figure_4.R')

cat("Creating Figure 5 Panels\n")
source('figure_5.R')

cat("Creating Supplemental Figure 5 Panels\n")
source('supplemental_figure_5.R')

cat("Creating Figure 6 Panels\n")
source('figure_6.R')

cat("Creating Figure 7 Panels\n")
source('figure_7.R')

cat("Figures for Jackson et al have been created in: ", file.path(WORKING.PATH, OUT.PATH), "\n")
quit(save = "no")
```
