## Supplementary material for "Gene regulatory network reconstruction using single-cell RNA sequencing of barcoded genotypes in diverse environments": Jackson_2019_Supplemental_Data_1.tar.gz: README

## Supplemental Data 1: Jackson et al 2019 ##
# Included in this archive are the following data files:
# data/
#	103118_SS_Data.tsv.gz (TSV count matrix of single-cell yeast reads [cells x genes])
#	110518_SS_NEG_Data (TSV count matrix of simulated single-cell reads [cells x genes])
#	yeast_gene_names.tsv (TSV with Systematic Names and Common Names for yeast genes)
#	STable5.tsv (TSV copy of Supplemental Table 5 with cross-validation results)
#	go_slim_mapping.tab (GO Slim Terms: https://downloads.yeastgenome.org/curation/literature/go_slim_mapping.tab)
# network/
#	signed_network.tsv (TSV signed [-1, 0, 1] network of regulatory relationships [genes x TFs])
#	COND_signed_network.tsv (TSV signed [-1, 0, 1] network of regulatory relationships [genes x TFs] for each of 11 conditions)
# priors/
#	Tchourine_gold_standard.tsv.gz (TSV with gold standard from Tchourine et al 2018)
#	ATAC-motif_priors.tsv.gz (TSV with atac-motif priors from Castro et al 2019)
#	YEASTRACT_priors_20181118.tsv.gz (TSV with YEASTRACT priors downloaded from YEASTRACT 11/18/2018)
#	BUSSEMAKER_priors_2008.tsv.gz (TSV with priors from Ward & Bussemaker 2008)

## To generate the figures in Jackson et al:

# 1) Edit jackson_2019_figures.R to change path or file names IF NEEDED.
# 2) Install the following packages (Note that this README can be run as an R script for installation)
# 3) Run the jackson_2019_figures.R script from the command line

# Install UMAP python package (the R UMAP library is a wrapper for this) with the appropriate manager:
# PyPI installation
# pip install umap-learn
# Conda installation
# conda install -c conda-forge umap-learn 

## Install required R packages ##
## Tidyverse ##
install.packages("bindrcpp", repo="http://cran.rstudio.com/")
install.packages("dplyr", repo="http://cran.rstudio.com/")
install.packages("reshape2", repo="http://cran.rstudio.com/")
install.packages("stringr", repo="http://cran.rstudio.com/")
install.packages("data.table", repo="http://cran.rstudio.com/")

## Plotting ##
install.packages("umap", repo="http://cran.rstudio.com/")
install.packages("igraph", repo="http://cran.rstudio.com/")
install.packages("scales", repo="http://cran.rstudio.com/")
install.packages("viridis", repo="http://cran.rstudio.com/")
install.packages("ggplot2", repo="http://cran.rstudio.com/")
install.packages("ggridges", repo="http://cran.rstudio.com/")
install.packages("cowplot", repo="http://cran.rstudio.com/")
install.packages("RColorBrewer", repo="http://cran.rstudio.com/")

## Math ##
install.packages("corpcor", repo="http://cran.rstudio.com/")
install.packages("irlba", repo="http://cran.rstudio.com/")

## BioConductor single-cell packages & DESeq2 ##
install.packages("BiocManager", repo="http://cran.rstudio.com/")
BiocManager::install(c("scater", "scran"))
BiocManager::install("DESeq2")

## Run the figure script ##
# Rscript jackson_2019_figures.R

## If you are particularly brave (or foolhearty), you may also wish to perform
## data imputation for network inference, or to generate Supplemental Figure 5
## Troubleshooting the following is left as an exercise to the reader. Beware.

## VIPER ##
install.packages("devtools", repo="http://cran.rstudio.com/")
install.packages("quadprog", repo="http://cran.rstudio.com/")
install.packages("glmnet", repo="http://cran.rstudio.com/")
library(devtools)
install_github("ChenMengjie/VIPER")

## SCIMPUTE ##
# install.packages("devtools", repo="http://cran.rstudio.com/")
library(devtools)
install_github("Vivianstats/scImpute")

## MAGIC ##
install.packages("Rmagic", repo="http://cran.rstudio.com/")
# MAGIC also requires a python package
# PyPI installation
# pip install --user magic-impute
# Conda installation
# conda install -c hcc magic

## Run the imputation script ##
# Rscript data_imputation.R
