## Supplementary material for "Gene regulatory network reconstruction using single-cell RNA sequencing of barcoded genotypes in diverse environments": Jackson_2019_Supplemental_Data_1.tar.gz: supplemental_figure_2.R

```
# Output paths
SUPPLEMENTAL.FIGURE.2.FILE <- "supplemental_figure_2.pdf"
NUM.CELLS.OUT <- "supplemental_figure_2a.png"
UMI.OUT <- "supplemental_figure_2b.png"
UMI.COND.OUT <- "supplemental_figure_2c.png"
UMI.GENO.OUT <- "supplemental_figure_2d.png"
CV.COND.OUT <- "supplemental_figure_2e.png"

if (!exists('count_data')) {source('figure_common_functions.R')}

count_data %>%
  dplyr::select(-c('Replicate', 'tenXBarcode')) %>%
  dplyr::mutate(umi=rowSums(.[1:6829])) %>%
  dplyr::group_by(Condition, Genotype_Group, Genotype) %>%
  dplyr::summarise(NumCells = n(), TotalUMI = sum(umi)) %>%
  dplyr::mutate(MeanUMI = TotalUMI / NumCells) -> df_summary_cells

plotter <- function(data, aes_for_data) {
  ggplot(data, aes_for_data) +
    theme_classic() +
    theme(axis.text.x = element_text(size = 8, angle=90, vjust=0.5), axis.title.x = element_text(size = 12),
          axis.text.y = element_text(size = 10), axis.title.y = element_text(size = 12),
          legend.text = element_text(size = 10), legend.title = element_text(size = 12, face = 'bold'),
          plot.title = element_text(size = 12, face = "bold", hjust=0.5),
          legend.position = "none")
}

pl1 <- plotter(df_summary_cells, aes(Genotype_Group, NumCells)) +
  geom_point(aes(color=Genotype_Group)) +
  scale_color_manual(labels = GENOTYPE.LABELS, breaks = GENOTYPE.LEVELS, values = GENOTYPE.COLORS) +
  scale_x_discrete(labels = rev(GENOTYPE.LABELS), limits = rev(GENOTYPE.LEVELS)) +
  facet_wrap(~Condition, labeller = as_labeller(structure(CONDITION.LABELS, names=CONDITION.LEVELS)), ncol = 6, scales="free") +
  labs(title="Unique Cells Per Biological Replicate", x="Genotype", y="Number of Cells", color="Genotype") +
  stat_summary(fun.y='mean', size=10, geom='point', shape='-') +
  theme(strip.background = element_blank(), strip.text = element_text(size=10, face = "bold"))

#ggsave(NUM.CELLS.OUT, plot=pl1, path=OUT.PATH, device="png", width=8.5, height=6, units="in")

pl2 <- plotter(df_summary_cells, aes(Genotype_Group, MeanUMI)) +
  geom_point(aes(color=Genotype_Group)) +
  scale_color_manual(labels = GENOTYPE.LABELS, breaks = GENOTYPE.LEVELS, values = GENOTYPE.COLORS) +
  scale_x_discrete(labels = rev(GENOTYPE.LABELS), limits = rev(GENOTYPE.LEVELS)) +
  facet_wrap(~Condition, labeller = as_labeller(structure(CONDITION.LABELS, names=CONDITION.LEVELS)), ncol = 6, scales="free_x") +
  labs(title="Mean UMI Per Biological Replicate", x="Genotype", y="UMI Count", color="Genotype") +
  stat_summary(fun.y='mean', size=10, geom='point', shape='-') +
  theme(strip.background = element_blank(), strip.text = element_text(size=10, face = "bold"))

#ggsave(UMI.OUT, plot=pl2, path=OUT.PATH, device="png", width=8.5, height=6, units="in")

count_data %>%
  dplyr::mutate(umi=rowSums(.[1:6829])) %>%
  dplyr::select(c('Condition', 'Genotype_Group', 'umi')) -> df_summary

pl3 <- plotter(df_summary, aes(Condition, umi)) +
  scale_fill_manual(labels = CONDITION.LABELS, breaks = CONDITION.LEVELS, values = CONDITION.COLORS) +
  scale_x_discrete(labels = rev(CONDITION.LABELS), limits = rev(CONDITION.LEVELS)) +
  geom_violin(aes(fill=Condition), scale = 'width') +
  geom_boxplot(outlier.shape = NA, width=0.25) +
  scale_y_continuous(limits=c(0,10000)) +
  labs(title="UMI Counts per Cell per Condition", x="Condition", y="UMI Count")

#ggsave(UMI.COND.OUT, plot=pl3, path=OUT.PATH, device="png", width=4, height=4, units="in")

pl4 <- plotter(df_summary, aes(Genotype_Group, umi)) +
  geom_violin(aes(fill=Genotype_Group), scale = 'width') +
  scale_fill_manual(labels = GENOTYPE.LABELS, breaks = GENOTYPE.LEVELS, values = GENOTYPE.COLORS) +
  scale_x_discrete(labels = rev(GENOTYPE.LABELS), limits = rev(GENOTYPE.LEVELS)) +
  geom_boxplot(outlier.shape = NA, width=0.25) +
  scale_y_continuous(limits=c(0,10000)) +
  labs(title="UMI Counts per Cell per Genotype", x="Genotype", y="UMI Count")

#ggsave(UMI.GENO.OUT, plot=pl4, path=OUT.PATH, device="png", width=4, height=4, units="in")

supp.plt.2 <- plot_grid(pl1, pl2, plot_grid(pl3, pl4, align="h", labels=c("C)", "D)")), ncol = 1, labels = c("A)", "B)", ""), rel_heights = c(1, 1, 0.75))
ggsave(SUPPLEMENTAL.FIGURE.2.FILE, plot=supp.plt.2, path=OUT.PATH, device=cairo_pdf, width=8.5, height=11, units="in", dpi = FIGURE.DPI)
```
