## Supplementary material for "Gene regulatory network reconstruction using single-cell RNA sequencing of barcoded genotypes in diverse environments": Jackson_2019_Supplemental_Data_1.tar.gz: supplemental_figure_4.R

```
# Output paths
SUPPLEMENTAL.FIGURE.4.FILE <- 'supplemental_figure_4.pdf'
SUPPLEMENTAL.FIGURE.4C.NETWORK.FILE <- 'supplemental_figure_4c_network_file.tsv'

if (!exists('count_data')) {source('figure_common_functions.R')}
if (!exists('sce.data')) {sce.data <<- dimred_for_umap(load_data_into_sce(count.data = dplyr::select(count_data, -META.DATA.COLUMNS), 
                                                                          meta.data = meta_data), 
                                                       random_state=RANDOM.SEED)}

print("Creating Supplemental Figure 4A")

sfig.4a.conditions <- c("YPD", "YPDRapa", "AmmoniumSulfate", "Urea")
sfigure.4a.mep2 <- plt.gene.violins(sce.data, "YNL142W", conditions = sfig.4a.conditions) + 
  facet_wrap(~condition, labeller = as_labeller(structure(CONDITION.LABELS, names=CONDITION.LEVELS)), ncol = 6) +
  labs(title="MEP2 Expression", x="Genotype", y="log2(Expression)")

sfigure.4a.gln1 <- plt.gene.violins(sce.data, "YPR035W", conditions = sfig.4a.conditions) + 
  facet_wrap(~condition, labeller = as_labeller(structure(CONDITION.LABELS, names=CONDITION.LEVELS)), ncol = 6) +
  labs(title="GLN1 Expression", x="Genotype", y="log2(Expression)")

sfigure.4a.hxk2 <- plt.gene.violins(sce.data, "YGL253W", conditions = sfig.4a.conditions) + 
  facet_wrap(~condition, labeller = as_labeller(structure(CONDITION.LABELS, names=CONDITION.LEVELS)), ncol = 6) +
  labs(title="HXK2 Expression", x="Genotype", y="log2(Expression)")

sfigure.4a <- plot_grid(sfigure.4a.mep2, sfigure.4a.gln1, sfigure.4a.hxk2, ncol = 1, align = "lr", labels = c("i", "ii", "iii"))

#ggsave(SUPPLEMENTAL.FIGURE.4A.FILE, plot=sfigure.4a, path=OUT.PATH, device="png", width=8.5, height=7.5, units="in")

print("Creating Supplemental Figure 4B")
non.wt.genotypes <- GENOTYPE.LEVELS[GENOTYPE.LEVELS != "WT(ho)"]
non.wt.genotype.labels <- GENOTYPE.LABELS[GENOTYPE.LABELS != "WT"]

markers.vs.wt <- NULL
for (condition  in CONDITION.LEVELS) {
  # subset sce for particular condition
  sce.sub <- sce.data[, sce.data$Condition == condition]
  # run pairwise T tests for all genotypes
  markers.out <- scran::pairwiseTTests(SummarizedExperiment::assay(sce.sub, 'logcounts'), sce.sub$Genotype_Group, lfc=FIG4.LFC.THRESHOLD) #
  # get pairs in which mutant is compared to wt
  pairs <- as.data.frame(markers.out$pairs)
  
  for (genotype in non.wt.genotypes) {
    geno.markers <- as.data.frame(markers.out$statistics[pairs$first == genotype & pairs$second == 'WT(ho)'])
    geno.markers$gene <- rownames(geno.markers)
    rownames(geno.markers) <- NULL
    geno.markers <- geno.markers[geno.markers$FDR < FIG4.P.ADJ.THRESHOLD, ]
    if (base::nrow(geno.markers) == 0) {next}
    geno.markers$genotype <- genotype
    geno.markers$condition <- condition
    markers.vs.wt <- rbind(markers.vs.wt, geno.markers)
  }
}

markers.vs.condition <- NULL
conditions.tested <- CONDITION.LEVELS
# run pairwise T tests for all conditions
markers.out <- scran::pairwiseTTests(assay(sce.data, 'logcounts'), sce.data$Condition, lfc=FIG4.LFC.THRESHOLD)
pairs <- as.data.frame(markers.out$pairs)

while (length(conditions.tested) > 1) {
  first.condition <- conditions.tested[1]
  conditions.tested <- conditions.tested[2:length(conditions.tested)]
  for (second.condition in conditions.tested) {
    markers.stats <- as.data.frame(markers.out$statistics[pairs$first == first.condition & pairs$second == second.condition])
    markers.stats$gene <- rownames(markers.stats)
    rownames(markers.stats) <- NULL
    markers.stats <- markers.stats[markers.stats$FDR < FIG4.P.ADJ.THRESHOLD, ]
    if(base::nrow(markers.stats) == 0) {next;}
    markers.stats$condition1 <- first.condition
    markers.stats$condition2 <- second.condition
    markers.vs.condition <- rbind(markers.vs.condition, markers.stats)
  }
}

# Summarize the differential gene expression results for plotting
markers.vs.wt %>%
  dplyr::group_by(condition, genotype) %>%
  dplyr::summarise(DiffGenes = n()) %>%
  dplyr::right_join(base::expand.grid(condition = CONDITION.LEVELS, 
                                      genotype = non.wt.genotypes, 
                                      stringsAsFactors = FALSE), 
                    by=c("condition", "genotype")) %>%
  as.data.frame() -> supp.figure.4b1.data

supp.figure.4b1.plt <- ggplot(supp.figure.4b1.data, aes(x=condition, y=genotype, fill=DiffGenes)) +
  theme_classic() +
  geom_tile() +
  geom_text(aes(label=DiffGenes)) +
  scale_x_discrete(limits=rev(CONDITION.LEVELS), labels=rev(CONDITION.LABELS)) +
  scale_y_discrete(limits=non.wt.genotypes, labels=non.wt.genotype.labels) +
  scale_fill_gradient(low="white", high=muted("blue"), na.value = "white") +
  theme(axis.text.x = element_text(size = 10, angle = 90, vjust=0.5, hjust=1), axis.title.x = element_text(size=12),
        axis.text.y = element_text(size = 10), axis.title.y = element_text(size=12),
        plot.title = element_text(size = 12, face = "bold", hjust = 0.5),
        legend.position = "none") +
  labs(x = "Growth Condition", y = "Genotype", title = "Differential Expression vs WT [# Genes]")

markers.vs.condition %>%
  dplyr::group_by(condition1, condition2) %>%
  dplyr::summarise(DiffGenes = n()) %>%
  dplyr::right_join(base::expand.grid(condition1 = CONDITION.LEVELS, 
                                      condition2 = CONDITION.LEVELS, 
                                      stringsAsFactors = FALSE), 
                    by=c("condition1", "condition2")) %>%
  as.data.frame() -> supp.figure.4b2.data

supp.figure.4b2.plt <- ggplot(supp.figure.4b2.data, aes(x=condition2, y=condition1, fill=DiffGenes)) +
  theme_classic() +
  geom_tile() +
  geom_text(aes(label=DiffGenes)) +
  scale_x_discrete(limits=rev(CONDITION.LEVELS), labels=rev(CONDITION.LABELS)) +
  scale_y_discrete(limits=CONDITION.LEVELS, labels=CONDITION.LABELS) +
  scale_fill_gradient(low="white", high=muted("blue"), na.value = "white") +
  theme(axis.text.x = element_text(size = 10, angle = 90, vjust=0.5, hjust=1), axis.title.x = element_text(size=12),
        axis.text.y = element_text(size = 10), axis.title.y = element_text(size=12),
        plot.title = element_text(size = 12, face = "bold", hjust = 0.5),
        legend.position = "none") +
  labs(x = "Growth Condition", y = "Growth Condition", title = "Differential Expression [# Genes]")

sfigure.4b <- plot_grid(supp.figure.4b1.plt, supp.figure.4b2.plt, align = "h", ncol = 2, labels = c("i", "ii"))
sfigure.4 <- plot_grid(sfigure.4a, sfigure.4b, align = "v", ncol=1, labels = c("A)", "B)"), rel_heights = c(1, 0.6))

ggsave(SUPPLEMENTAL.FIGURE.4.FILE, plot=sfigure.4, path=OUT.PATH, device=cairo_pdf, width=8.5, height=10, units="in", dpi = FIGURE.DPI)
```
