## Supplementary material for "Gene regulatory network reconstruction using single-cell RNA sequencing of barcoded genotypes in diverse environments": Jackson_2019_Supplemental_Data_1.tar.gz: supplemental_figure_5.R

```
# Output paths
SUPP.FIG.5 <- "supplemental_figure_5a.png"

if (!exists('count_data')) {source('figure_common_functions.R')}

if (!exists('IMPUTED.DATA.FILES')) {IMPUTED.DATA.FILES <<- list(Unmodified=list("103118_SS_Data.tsv.gz", TRUE),
                                                                MAGIC=list("MAGIC_DATA.tsv.gz", FALSE),
                                                                scImpute=list("SCIMPUTE_DATA.tsv.gz", FALSE),
                                                                VIPER=list("VIPER_DATA.tsv.gz", TRUE))}

load_data <- function(data.file.name, genes.are.columns=TRUE, meta.data.columns=NULL) {
  loaded_count_data <- as.data.frame(fread(file.path(WORKING.PATH, DATA.PATH, data.file.name)))
  rownames(loaded_count_data) <- loaded_count_data[, 1]
  loaded_count_data <- loaded_count_data[, -1]
  loaded_count_data <- fix_imported_data_frame(loaded_count_data)
  
  if (!is.null(meta.data.columns)) {
    loaded_meta_data <- dplyr::select(loaded_count_data, meta.data.columns)
    loaded_count_data <- dplyr::select(loaded_count_data, -meta.data.columns)
    sce.obj <- load_data_into_sce(loaded_count_data, loaded_meta_data, genes.are.columns = genes.are.columns)
  }
  else {
    sce.obj <- load_data_into_sce(loaded_count_data, meta_data, genes.are.columns = genes.are.columns)
  }
  
  return (sce.obj)
}

plt.list = list()
for (imputed.method in names(IMPUTED.DATA.FILES)) {
  # Check to see if the data file exists
  if (!file.exists(file.path(WORKING.PATH, DATA.PATH, IMPUTED.DATA.FILES[[imputed.method]][[1]]))) {
    print(paste0("Skipping data from ", imputed.method, " (File Not Found)")) 
    next
  }
  
  # Load and process count data file
  print(paste0("Loading data: ", imputed.method))
  sce.obj <- load_data(IMPUTED.DATA.FILES[[imputed.method]][[1]], meta.data.columns = META.DATA.COLUMNS)
  sce.obj <- dimred_for_umap(sce.obj, do.multibatchnorm = IMPUTED.DATA.FILES[[imputed.method]][[2]])
  
  print(paste0("Plotting UMAP: ", imputed.method))
  p1 <- umap.plot(sce.obj, color.by='Condition', alpha=0.5, size=0.05, color.title='Condition') + 
    scale_color_manual(labels = CONDITION.LABELS, breaks = CONDITION.LEVELS, values = CONDITION.COLORS) + 
    guides(colour = guide_legend(override.aes = list(size=4, alpha=1), reverse = TRUE)) +
    labs(title = paste0("Growth Condition\n[", imputed.method,"]")) +
    theme(legend.position = "none", axis.title = element_blank(), axis.text = element_blank(), 
          plot.title = element_text(size=10, face="bold", hjust=0.5))

  p2 <- umap.plot(sce.obj, color.by='Genotype_Group', alpha=0.5, size=0.05, color.title='Genotype') + 
    scale_color_manual(labels = GENOTYPE.LABELS, breaks = GENOTYPE.LEVELS, values = GENOTYPE.COLORS) + 
    guides(colour = guide_legend(override.aes = list(size=4, alpha=1), reverse = TRUE)) +
    labs(title = paste0("Strain Genotype\n[", imputed.method,"]")) +
    theme(legend.position = "none", axis.title = element_blank(), axis.text = element_blank(), 
          plot.title = element_text(size=10, face="bold", hjust=0.5))
  
  plt.combined <- plot_grid(p1, p2, ncol = 2, align = 'v')
  plt.list[[imputed.method]] <- plt.combined
  #ggsave(plot=plt.combined, filename=paste0(substr(SUPP.FIG.5, 1, 22), "_", imputed.method, ".png"), path=OUT.PATH, device="png", width=8.5, height=4, units="in")
}

plt.all <- plot_grid(plotlist = plt.list, ncol = 2, align = "hv", labels = c("A)", "", "", ""))
ggsave(SUPP.FIG.5, plot = plt.all, path = OUT.PATH, width = 8.5, height = 5, device = "png", units = "in", dpi = FIGURE.DPI)
```
